# Tonotopic Specialization of MYO7A Isoforms in Auditory Hair Cells

**DOI:** 10.1101/2025.04.01.646665

**Authors:** Sihan Li, Jinho Park, Tobey M. Phan, Edward H. Egelman, Jonathan E. Bird, Jung-Bum Shin

## Abstract

Mutations in *Myo7a* cause Usher syndrome type 1B and non-syndromic deafness, but the precise function of MYO7A in sensory hair cells remains unclear. We identify and characterize a novel isoform, MYO7A-N, expressed in auditory hair cells alongside the canonical MYO7A-C. Isoform-specific knock-in mice reveal that inner hair cells primarily express MYO7A-C, while outer hair cells express both isoforms in opposing tonotopic gradients. Both localize to the upper tip-link insertion site, consistent with a role in the tip link for mechanotransduction. Loss of MYO7A-N leads to outer hair cell degeneration and progressive hearing loss. Cryo-EM structures reveal isoform-specific differences at actomyosin interfaces, correlating with distinct ATPase activities. These findings reveal an unexpected layer of molecular diversity within the mechanotransduction machinery. We propose that MYO7A isoform specialization enables fine-tuning of tip-link tension, thus hearing sensitivity, and contributes to the frequency-resolving power of the cochlea.

## 2. Introduction

Precise detection of sound relies on the exquisite mechanical sensitivity of sensory hair cells located in the inner ear^1,2^. These specialized sensory receptors convert minute mechanical vibrations into electrical signals through a process known as mechano-electrical transduction (MET)^1–3^. At the heart of this process lies the hair bundle—a staircase-like array of actin-based stereocilia^4^—whose deflection modulates tension on filamentous structures called tip links, ultimately gating MET channels^1–3^.

The MET complex is positioned between adjacent rows of stereocilia, with channels located at the tips of shorter stereocilia and tethered to the taller neighbors via tip links^5,6^ (**Fig. 1A**). Deflection of the hair bundle increases tip-link tension, opening MET channels and initiating receptor potentials.

**Figure 1:**
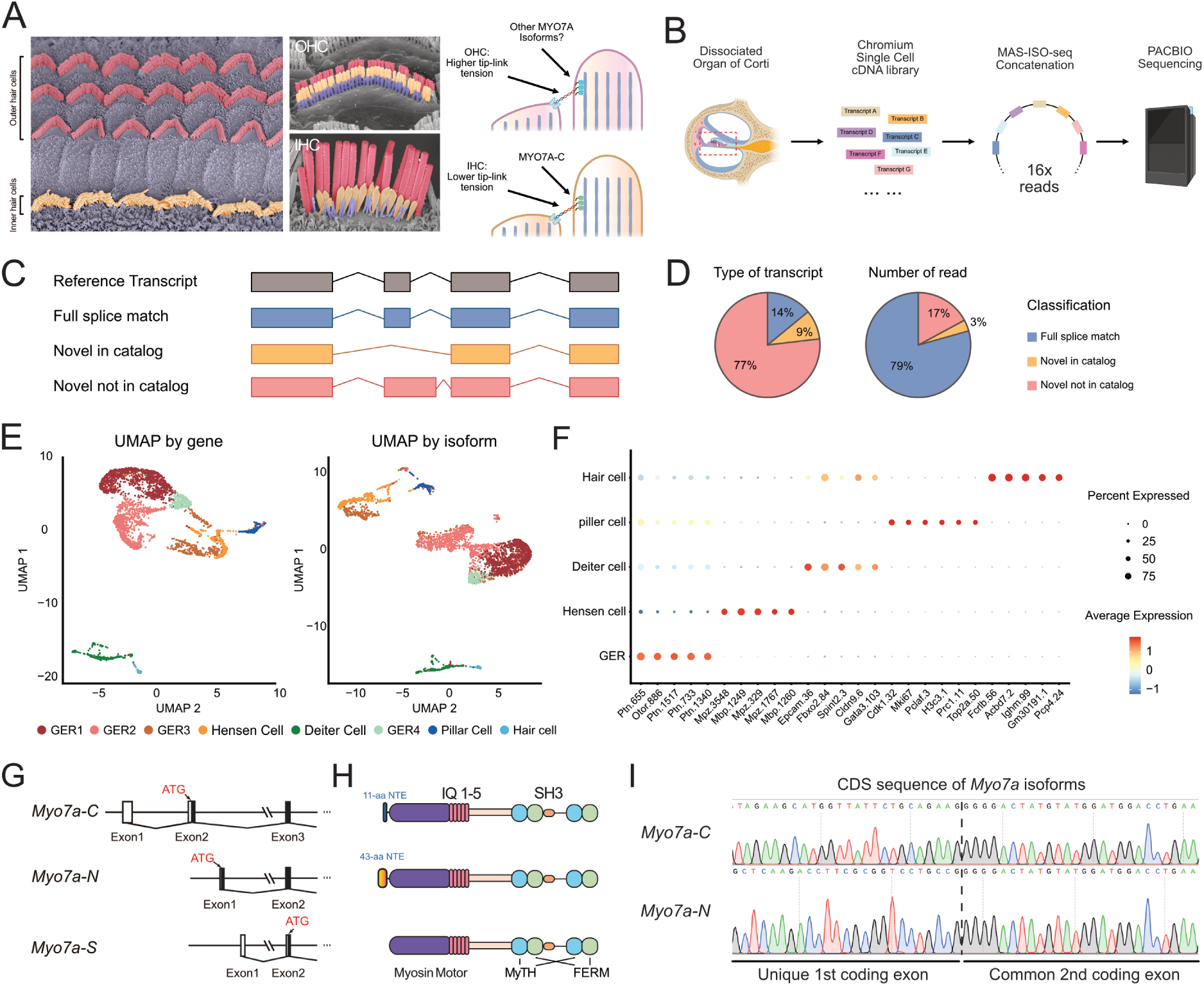
Identification of MYO7A-N by PACBIO MAS-ISO-Seq. **A.** SEM images of the organ of Corti, and schematic illustration of tip link complex. **B**. Workflow of MAS-ISO-seq. 10x Genomics 3’ Single cell cDNA library was generated from dissociated Organ of Corti. Then the cDNA was processed by PACBIO MAS-ISO-seq kit to concatenated sixteen transcripts into each SMRTbell. Lastly, the SMRTbell library was sequenced by PACBIO Revio system. **C**. Overview of RNA transcript category according to Iso-Seq analysis, including full splice match, novel in catalog, novel not in catalog. **D**. Pie chart demonstrating the percentage of RNA isoforms by type of transcript category or number of read detected. **E**. Uniform manifold approximation and projection (UMAP) plot of single cells colored by cell types by gene or isoform expression. All the cells are classified into 8 clusters. **F**. Dot plot heatmap of average expression and cellular detection rate for top 5 isoforms for each cell type. Different GER cell types are grouped into one group. **G**. Diagram of the 5’ end of *Myo7a* isoforms identified by MAS-ISO-Seq in cochlear hair cells. *Myo7a* is expressed as three different isoforms: *Myo7a-C* and *Myo7a-S* are reported in genomic databases. The newly identified *Myo7a-N* isoform utilizes a unique, previously unreported exon. These three isoforms are generated by different transcriptional start sites and translational start sites (ATG). **H**. Illustration of protein domains of MYO7A isoforms. Compared to MYO7A-S, MYO7A-C has an 11-aa extension at the N-terminus. The newly discovered isoform of MYO7A-N has a 43-aa N-terminus extension. **I**. Sanger sequencing result of *Myo7a-C* and *Myo7a-N* transcripts from 5’ RACE cDNA library, confirming *Myo7a-C* and *Myo7a-N* have unique first coding exons.

Many core components of this complex have been identified: TMC1 and TMC2 form the MET channel pore^7–11^, while auxiliary proteins such as TMIE^7,12,13^, LHFPL5^13–15^, CIB2, and CIB3^16,17^ and others are essential for proper channel function. At the upper insertion point of the tip link—the upper tip-link density (UTLD)—a scaffold complex anchors the tip link to the actin core and helps maintain appropriate tension^5,18^. Proper tip-link tension is essential not only for fast and precise MET channel gating, but also for tuning the overall sensitivity of hair cells to mechanical stimuli—thereby directly impacting hearing sensitivity^5,19,20^. Reduced tip-link tension delays channel opening and diminishes the receptor potential amplitude, compromising the fidelity of sound encoding^20^.

Interestingly, tip-link tension is not uniform across hair cell types or cochlear regions. Outer hair cells (OHCs) are thought to have higher tip-link tension than inner hair cells (IHCs), and within OHCs, tension increases along the tonotopic axis from apex to base^19^. These gradients suggest that hair cells dynamically optimize tip-link tension to match their frequency tuning properties and preserve auditory acuity.

Proteins localized to the UTLD include Harmonin (USH1C), Sans (USH1G), and the motor protein Myosin VIIA (MYO7A)^18^. Harmonin and Sans serve as scaffolds, while MYO7A is proposed to actively generate or stabilize tip-link tension through interactions with actin filaments^18^. Notably, *MYO7A* is a well-established human deafness gene and the causal gene for Usher syndrome type 1B, a disorder characterized by congenital deafness, vestibular areflexia, and progressive vision loss^21–23^.

Our previous work demonstrated that deletion of the canonical MYO7A isoform (MYO7A-C) markedly reduced MYO7A protein in IHCs, decreased MET resting open probability, and slowed MET current activation, consistent MYO7A’s proposed role in maintaining tip-link tension^20^. However, this functional role remains under debate. A recent study proposed that MYO7A may act primarily as a structural anchor, passively linking the tip link to the actin cytoskeleton rather than actively generating tension^24^. In this model, tip-link tension could instead arise from alternative mechanisms, such as actin remodeling at the transducing row of stereocilia or tension in the membrane^25^. Thus, the precise role of MYO7A in regulating tip-link tension—and whether it functions as a motor, scaffold, or both—remains to be resolved.

In this study, we used long-read RNA sequencing to identify a novel MYO7A isoform, MYO7A-N, expressed in auditory hair cells (**Fig. 1B**). MYO7A-N is predominantly expressed in OHCs, following a basal-to-apical tonotopic gradient, whereas MYO7A-C is enriched in IHCs and displays an opposing gradient in OHCs. Notably, both isoforms are localized at the UTLD, where tip-link tension is regulated. Functional deletion of MYO7A-N in mice leads to progressive hearing loss and OHC degeneration. Cryo-EM analysis reveals structural divergence in their motor domains, corresponding to distinct ATPase activities. These findings uncover isoform-specific roles of MYO7A in shaping tip-link tension and suggest a novel mechanism for fine-tuning MET sensitivity and auditory function across hair cell populations.

## 3. Results

### 3.1 Survey of cochlear *Myo7a* isoforms by long-read sequencing

Two major types of *Myo7a* isoforms have been identified in multiple genomic databases: the canonical isoform (*Myo7a-C*) and the short isoform (*Myo7a-S*). These isoforms differ in their transcriptional and translational start sites (ATG), resulting in distinct N-terminal extensions (NTEs) of the motor domain. Our previous study demonstrated that *Myo7a-C* is predominantly expressed in all cochlear IHCs and exhibits a tonotopic gradient in OHCs^20^. However, the existence and function of *Myo7a-S*, along with the potential presence of other isoforms is unknown.

To comprehensively characterize the isoform transcript landscape in cochlear cells, including *Myo7a* isoforms featuring alternative NTEs, we employed multiplexed arrays isoform sequencing (MAS-ISO-seq) on cochleae from postnatal day 7 (P7) mice^26^ (**Fig. 1B**). This single-cell long-read sequencing technique generates a detailed catalog of isoforms across different cochlear cell types (**Fig. 1C**). Consistent with previous studies using a different long-read sequencing approach in cochlea^27^, we identified numerous previously unreported isoforms. After filtering ambiguous transcripts such as intergenic transcripts and incomplete transcripts, 26,314,245 (79%) transcripts fully matched the reference genome exon annotation (**Fig. 1D**). There are also 1,159,082 transcripts (3%) that were categorized as “novel in catalog,” indicating new isoforms using known splicing donor/acceptor sites in different combinations (**Fig. 1D**). An additional 5,681,939 transcripts (17%) were classified as “novel not in catalog”, representing alternative splicing events with previously unidentified donor/acceptor sites (**Fig. 1D**). As validation of this approach, we successfully identified both long and short isoforms of otoferlin (OTOF), consistent with a recent report^27^. Single cell RNAseq analysis pipeline identified a total of 3707 cells, which were classified into 8 distinct clusters, including hair cells, Hensen cells, Deiters’ cells and other cell types (**Fig. 1E**). Subsequent analysis detected distinctively expressed isoforms for each cell types (**Fig. 1F**).

Focusing on *Myo7a* in the hair cell cluster, we detected transcripts corresponding to the previously reported *Myo7a-C* and *Myo7a-S* isoforms. Interestingly, we also identified a novel isoform, which we named *Myo7a-N* (**Fig. 1G-I**). To validate the presence of *Myo7a-N*, we performed 5’ rapid amplification of cDNA ends (5’ RACE) on cochlear cDNA from P0 mice. *Myo7a-C* and *Myo7a-N* transcripts were successfully identified in the 5’ RACE cDNA library (**Fig. 1I**). However, *Myo7a-S* transcripts were not detected in the same library, suggesting that, if expressed, *Myo7a-S* is present at very low levels in cochlear hair cells.

Like *Myo7a-C* and *Myo7a-S*, *Myo7a-N* is generated from an alternative transcriptional start site. Sequencing revealed that the unique first exon of *Myo7a-N* is 145 bp long, with the first 31 nucleotides comprising an untranslated region (UTR), and the remaining sequence encoding a 36-amino-acid NTE. Our data thus reveals a spectrum of NTEs for MYO7A; the shortest isoform, MYO7A-S, lacks an NTE, whereas MYO7A-C has an 11 amino acid NTE, and MYO7A-N has a 36 amino acids NTE (**Fig. 1H**). Importantly, the 36-amino acid NTE of MYO7A-N is highly conserved across vertebrates, including mice and humans, indicating a potentially critical evolutionary role in hair cell function (**Supplementary Fig. 1A, B**).

### 3.2 MYO7A-C and MYO7A-N constitute the major MYO7A isoforms in the cochlea

While MAS-ISO-seq is a powerful tool in identifying isoform transcripts, the read depth was not sufficient for reliable quantification of transcripts. To quantify the transcript levels of the different *Myo7a* isoforms, we performed quantitative polymerase chain reaction (qPCR) from inner ear cDNA libraries. We isolated cochlear mRNA at P0 and P20 to trace expression level changes of *Myo7a* isoforms at different developmental stages. Consistent with our 5’ RACE result, *Myo7a-C* and *Myo7a-N* isoforms were the most abundant isoforms detected at P0. In contrast, the transcript of *Myo7a-S* was nearly undetectable (**Fig. 2A**).

**Figure 2:**
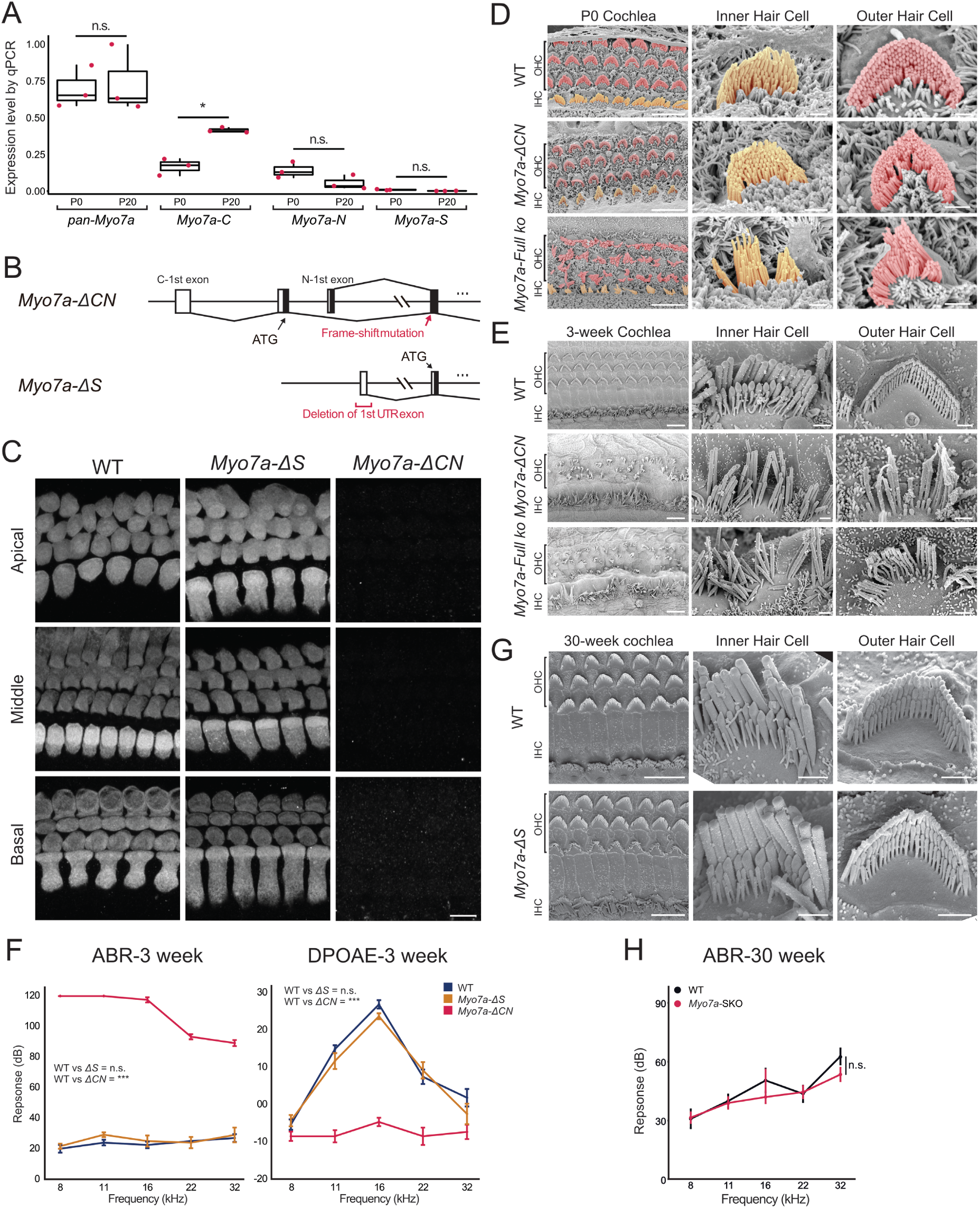
*Myo7a-C* and *Myo7a-N* are the major isoforms in cochlear hair cells. **A.** Normalized expression levels of *Myo7a* isoforms at P0 and P20 as determined by qPCR. Transcript levels were first normalized to *Gapdh* and then further normalized to the highest expressed transcript to calculate the relative expression levels. Compared to *Myo7a-C* and *Myo7a-N*, *Myo7a-S* has a low expression level in cochlea cDNA (each datapoint represents an individual cDNA preparation. Each cDNA preparation utilized 8∼9 cochleae. Normalized percentage: P0= 0.70, P20=0.74; p-value=0.82. T-test p-value of each isoform P0 vs. P20: *pan-Myo7a*=0.82, *Myo7a-C*=0.011, *Myo7a-N*=0.1162, *Myo7a-S*=0.060). Box-plots show medians, 25th, and 75th percentiles as box limits and minima and maxima as whiskers. **B**. Diagram describing the generation of *Myo7a-ΔCN* and *Myo7a-ΔS* mice. To generate the *Myo7a-ΔCN* mouse line, a single nucleotide deletion resulting in a frameshift was introduced in the coding sequence of both *Myo7a-C* and *Myo7a-N* but not *Myo7a-S*. To generate the *Myo7a-ΔS* mouse, the first non-coding first was deleted. **C**. Immunofluorescence images of wildtype (WT), *Myo7a-ΔS*, and *Myo7a-ΔCN* mice at P0. *Myo7a-ΔS* exhibits WT-like MYO7A immunofluorescent intensity in both IHCs and OHCs. In contrast, *Myo7a-ΔCN* mice showed a nearly abolished MYO7A signal, similar to *Myo7a-full KO* mice. **D**. SEM of WT, *Myo7a-ΔCN,* and *Myo7a full KO* mouse cochlea at P0. Despite a substantial reduction of MYO7A level, *Myo7a-ΔCN* cochlear hair cells retained WT-like hair bundle morphology at P0. *Myo7a-full KO* mice showed disorganized hair bundle morphology. Scale bar: view of the organ of Corti: 10µm; view of IHC or OHC: 1µm. **E**. SEM of WT, *Myo7a-ΔCN,* and *Myo7a* full KO mouse cochlea at P20. Hair bundle disorganization and hair cell loss is evident in *Myo7a-ΔCN* mice, comparable to *Myo7a-full KO* cochlea. Scale bar: view of the organ of Corti: 10µm; view of the IHC or OHC: 1µm. **F**. ABR and DPOAE test of WT, *Myo7a-ΔCN,* and *Myo7a-ΔS* mice at 3 weeks. *Myo7a-ΔCN* showed profound hearing loss with nearly abolished ABR and DPOAE. *Myo7a-ΔS* mice do not show a significant change in hearing performance. (ANOVA p-value: ABR: WT vs. *Myo7a-ΔCN*: 8.21e-38, WT vs. *Myo7a-ΔS*: 0.20. The number of animals: WT=6 vs. *Myo7a-ΔCN*=6, *Myo7a-ΔS*=6. DPOAE: WT vs *Myo7a-ΔCN*: 1.49e-12, WT vs *Myo7a-ΔS*: 0.19. Number of animals: WT=5 vs *Myo7a-ΔCN*=6, *Myo7a-ΔS*=6). Line-plots show the means of the ABR or DPOAE response. Error bars represent standard errors. **G**. SEM of WT and *Myo7a-ΔS* mouse cochlea at 30 weeks. The hair bundles of both IHCs and OHCs of *Myo7a-ΔS* mice retain a WT-like hair bundle structure. Scale bar: view of the organ of Corti: 10µm; view of IHC or OHC: 1µm. **H**. ABR and DPOAE test of WT and *Myo7a-ΔS* mice at 30 weeks. *Myo7a-ΔS* mice do not show a significant change in hearing performance. (ANOVA p-value: ABR: WT vs. *Myo7a-ΔCN*: 8.21e-38, WT vs. *Myo7a-ΔS*: 0.20. The number of animals: WT=6 vs. *Myo7a-ΔCN*=6, *Myo7a-ΔS*=6. DPOAE: WT vs *Myo7a-ΔCN*: 1.49e-12, WT vs *Myo7a-ΔS*: 0.19. Number of animals: WT=5 vs *Myo7a-ΔCN*=6, *Myo7a-ΔS*=6). Line-plots show the means of the ABR or DPOAE response. Error bars represent standard errors.

The expression levels of *Myo7a-C* and *Myo7a-N* changed during postnatal development between P0 and P20. The total level of *Myo7a* transcripts did not significantly change between P0 and P20, while the ratio of different isoforms changed during the course of hair cell maturation: the transcript level of *Myo7a-C* increased by approximately 2.5-fold from P0 to P20 (Normalized percentage: P0= 0.17, P20=0.42; p-value=0.011(*)). In contrast, the *Myo7a-N* trended towards a decrease (Normalized percentage: P0=0.14, P20=0.056; p-value=0.12). Finally, the transcript level of *Myo7a-S* was barely detectable and remained low during maturation (Normalized percentage: P0=0.0095, P20=0.0017; p-value=0.060). This result shows that *Myo7a-C* and *Myo7a-N* are the predominant isoforms in the cochlea, and that their expression levels are refined during cochlea maturation (**Fig. 2A**).

Although *Myo7a-S* is expressed at low levels, it is still possible that it has a specific and indispensable role in hair cell development or function. We tested this hypothesis using two mouse models. First, we generated a mouse line in which both *Myo7a-C* and *Myo7a-N* were deleted simultaneously. This mouse line was created by introducing a frameshift mutation in the common reading frame of the *Myo7a-C* and *Myo7a-N*, but upstream of the start codon of *Myo7a-S*. This allowed us to disrupt the translation of the MYO7A-C and MYO7A-N, without affecting MYO7A-S expression. We named this mouse line *Myo7a-ΔCN* mouse (**Fig. 2B**). In addition, we sought to generate a mouse line in which only *Myo7a-S* is disrupted. Because the start codon of the *Myo7a-S* isoform is located in the shared reading frame of the *Myo7a-C* and the *Myo7a-N* isoforms, it was not possible to interrupt the translated portion of *Myo7a-S* without affecting other isoforms. As an alternative method, we deleted the entire first exon that has the transcriptional start site of *Myo7a-S*, which theoretically should prevent the transcription *Myo7a-S*. This mouse line was named *Myo7a-ΔS* (**Fig. 2B**).

First, we characterized the expression pattern of MYO7A in the *Myo7a-ΔCN* and *Myo7a-ΔS* mouse lines using a pan-MYO7A antibody^20,28^. At P5, MYO7A immunofluorescence was abolished in all hair cells of *Myo7a-ΔCN* mice. In contrast, we did not observe a noticeable reduction of MYO7A immunoreactivity in *Myo7a-ΔS* cochlea hair cells. This result was consistent with our qPCR analysis and confirmed that MYO7A-C and MYO7A-N are the major expressed isoforms in cochlear hair cells (**Fig. 2C**).

Next, we used scanning electron microscopy (SEM) to examine the hair bundle morphology in *Myo7a-ΔCN* mice at P0 and P20 (**Fig. 2D-E**). Given the complete loss of MYO7A signal in P0 *Myo7a-ΔCN* hair cells, we anticipated a severely disorganized hair bundle phenotype^29^. Surprisingly, while both IHCs and OHCs exhibited morphological abnormalities, the disorganization was less severe than that observed in *Myo7a* full KO mice at P0 (**Fig. 2D**). This finding suggests that residual MYO7A isoforms—most likely MYO7A-S—can partially compensate for the absence of the major isoforms during embryonic hair bundle development. In contrast, by postnatal day 20, the hair bundle morphology in *Myo7a-ΔCN* mice had degenerated to a degree comparable to age-matched *Myo7a-full KO* mice (**Fig. 2E**). This morphological deterioration of the hair bundle suggests the level of remaining MYO7A (likely MYO7A-S) was not sufficient to maintain the hair bundle morphology in *Myo7a-ΔCN* mice. This result is consistent with our qPCR analysis that the expression level of *Myo7a-S* further decreases during hair cell maturation.

To test if MYO7A-S has a specific role in hair cell maintenance, we examined hair bundle morphology of aged *Myo7a-ΔS* mice (**Fig. 2G**). At 30-week, the hair bundle of *Myo7a-ΔS* mice appeared comparable to WT mice (**Fig. 2G**), suggesting that MYO7A-S is not necessary for developing or maintaining the hair bundle morphology in mature hair cells.

Finally, we assessed the relevance of MYO7A-S in hearing function by conducting auditory brainstem response (ABR) on *Myo7a-ΔS* and *Myo7a-ΔCN* mice at 3-weeks and 30-weeks. *Myo7a-ΔCN* mice suffered profound hearing loss and complete lack of distortion product otoacoustic emissions (DPOAE) at 3 weeks of age (**Fig. 2F**). This result is consistent with the disorganized hair bundle morphology shown by SEM (**Fig. 2E**). In contrast, *Myo7a-ΔS* mice did not show any significant difference in ABR threshold compared to age-matched WT controls at 3 weeks (**Fig. 2F**) and 30 weeks (**Fig. 2H**). Overall, we conclude that MYO7A-S is expressed at low levels during early development but is dispensable for hair bundle development and hearing. Consequently, we decided to focus on the functional differences between the MYO7A-C and the MYO7A-N in the remainder of our studies.

### 3.3 MYO7A-C and MYO7A-N exhibit opposing tonotopic gradients in the cochlea

To explore the function of MYO7A-N, we created a mouse line in which we specifically knocked out *Myo7a-N* by deleting its entire first coding exon (*Myo7a-ΔN* mouse) (**Fig. 3A**). Together with the *Myo7a-ΔC* mouse line we characterized in our previous study^20^, we were able to investigate the expression patterns and functional differences of MYO7A-C and MYO7A-N in the mouse cochlea. First, to check the expression pattern of these two isoforms, we performed immunohistochemistry experiments on *Myo7a-ΔC* and *Myo7a-ΔN* P5 cochleae using a pan-MYO7A antibody. As described in the previous study, MYO7A immunoreactivity in IHCs of *Myo7a-ΔC* mice is substantially decreased compared to WT^20^ (**Fig. 3B, C**). Homozygous *Myo7a-ΔC* mice also exhibited a tonotopic reduction of MYO7A signal in OHCs, with a stronger reduction at the apical region and a more modest reduction at the cochlear base (**Fig. 3B, C**). In contrast, homozygous *Myo7a-ΔN* mice showed a reduction in total MYO7A levels mainly in basal OHCs, and this reduction of MYO7A became less pronounced toward the apex of the cochlea. No significant MYO7A signal reduction was observed in *Myo7a-ΔN* IHCs. The quantification of total MYO7A immunofluorescence in the isoform-specific KO mice highlights and summarizes the inverse expression pattern of MYO7A-C and MYO7A-N: IHCs predominantly express MYO7A-C in all regions of the cochlea. OHCs are predominantly expressed in the OHCs with a tonotopic pattern, with a higher expression level at the basal region of the cochlea and decrease toward the apex (**Fig. 3D**).

**Figure 3:**
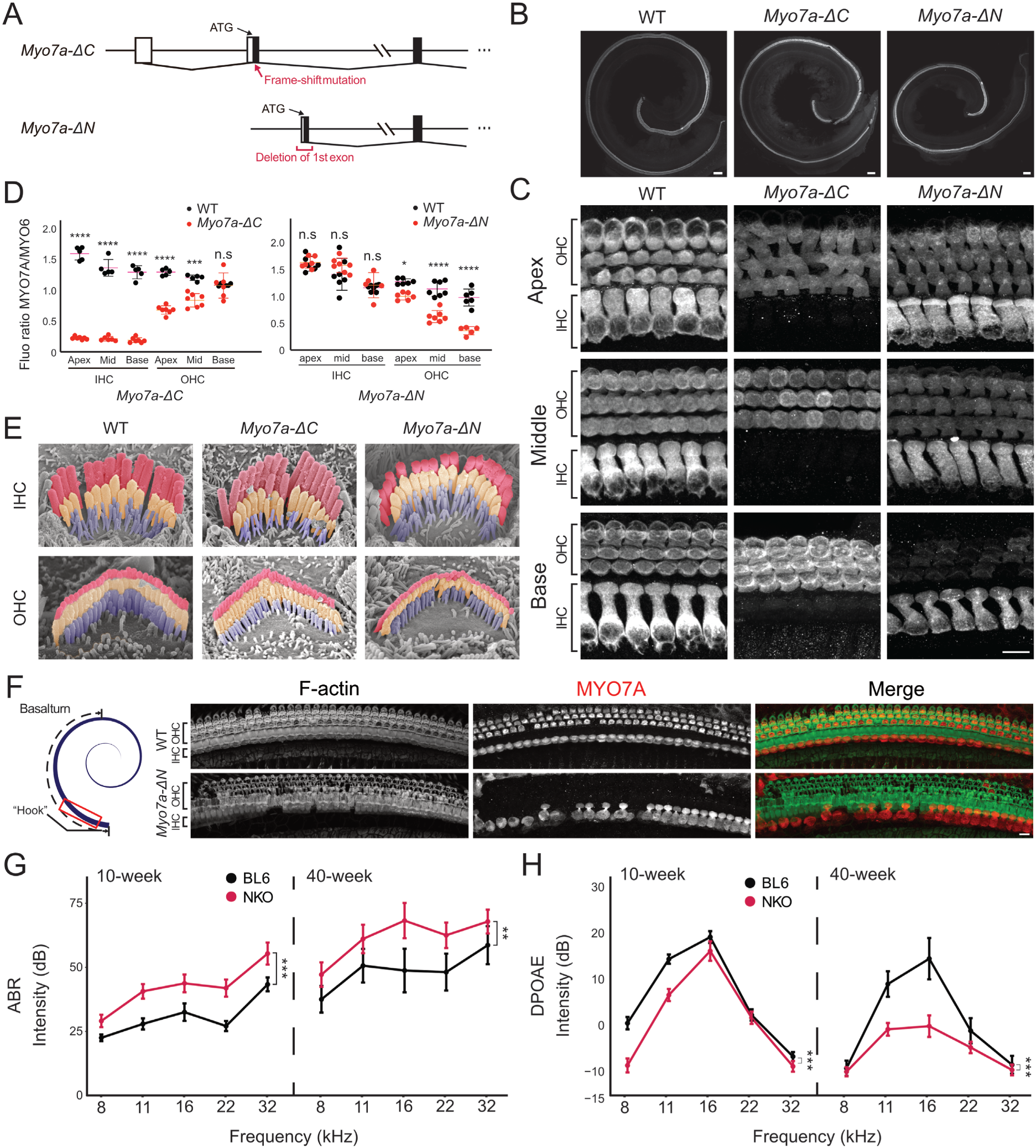
Expression of MYO7A-C and MYO7A-N are inversely correlated in the cochlea. **A.** Diagram of the generation of *Myo7a-ΔC* and *Myo7a-ΔN* mice. The entire first coding exon of *Myo7a-N* was deleted to generate the *Myo7a-ΔN* mice. Generation of *Myo7a-ΔC* mice was described in our previous study^20^. **B**. Pan-MYO7A immunofluorescence of organ of Corti whole mount from WT, *Myo7a-ΔC*, *Myo7a-ΔN* mice at P7. **C**. Zoomed-in images of WT, *Myo7a-ΔC*, *Myo7a-ΔN* ko mice at P7 at apical, middle, and basal region. Scale bar=10µm. **D**. Quantification of MYO7A immunofluorescence of WT, *Myo7a-ΔC*, *Myo7a-ΔN* mice in P7 cochlea. The quantification of fluorescence intensity showed that MYO7A-C and MYO7A-N are inversely correlated in cochlea hair cells. MYO7A immunoreactivity is normalized to MYO6 (not shown). Box-plots indicate medians, 25th, and 75th percentiles as box limits and minima and maxima as whiskers. **E**. SEM images of the IHC and OHC of WT, *Myo7a-ΔC*, and *Myo7a-ΔN* at P7. No morphological changes are detected in *Myo7a-ΔC*, and *Myo7a-ΔN* hair cells. **F**. Immunofluorescence images of WT and *Myo7a-ΔN* mice at 40 weeks. Images were taken at the beginning of the basal turn of the cochlea (“hook region”). The cochlea from *Myo7a-ΔN* mice showed substantial OHC loss. **G**, **H**. ABR and DPOAE of WT and *Myo7a-ΔN* mice at 10 and 40 weeks. Our results are consistent with deterioration of OHC function of *Myo7a-ΔN* with age (Number of animals: WT: 10-week: 9; 40-week: 8; *Myo7a-ΔN*: 10-week: 14, 40-week: 16. ANOVA p-value: 10-week: 3.80e-05; 40-week: 3.24e-05.)

### 3.4 Hair cell loss and progressive mild hearing loss in adult *Myo7a-ΔN* mice

It is well established that MYO7A is essential for hair bundle development, and that the depletion of MYO7A results in hair bundle disorganization^30,31^. We therefore investigated the morphology of hair cells and their hair bundles in *Myo7a-ΔN* mice by SEM imaging analysis. As reported in our previous study^20^, despite an approximately 80% reduction of MYO7A in IHCs, *Myo7a-ΔC* IHC hair bundles retain WT-like morphology at P7 (**Fig. 3E**). Similarly, *Myo7a-ΔN* basal OHCs, which show the most significant MYO7A reduction, did not show a substantial change in hair bundle morphology at young age. We then evaluated the effect of a specific deletion of *Myo7a-N* on hair cell maintenance at 40 weeks of age, by examining OHC bundles of the beginning region of the basal turn called the “hook” region. At 40-weeks of age, both IHCs and OHCs are well maintained in the WT mice. However, most of OHCs were lost in the basal turn of *Myo7a-ΔN* cochlea (**Fig. 3F**). This result showed that the deletion of MYO7A-N renders basal OHC more vulnerable to factors such as aging.

We also evaluated the hearing function of *Myo7a-ΔN* mice at 10 and 40 weeks. At both ages, ABR thresholds were significantly elevated and DPOAE responses were reduced (**Fig. 3G, H**). While the changes were significant, ABRs and DPOAEs were only mildly compromised in *Myo7a-ΔN* mice. This is consistent with the fact that deletion of MYO7A-N reduces total MYO7A levels in OHCs maximally by ∼60% even at basal regions. Nevertheless, these findings suggest that MYO7A-N is important for hearing function, likely related to its role in OHC function and cochlear amplification.

### 3.5 MYO7A-C and MYO7A-N localize to the UTLD in hair cells in a tonotopically graded manner

MYO7A at the UTLD is proposed to establish resting tension on the tip link by forming a complex with Harmonin and Sans, bridging the actin cytoskeleton with the cytosolic domains of CDH23.^18,32–34^. To test whether MYO7A isoforms exhibit distinct subcellular localizations, particularly at UTLD, we created mouse lines in which we inserted an HA-tag after the start codon of *Myo7a-C* and *Myo7a-N* (**Fig. 4A**). Using the *HA-Myo7a-N* and *HA-Myo7a-C* mouse lines, we visualized the expression and localization of the two isoforms separately at both tissue and subcellular levels.

**Figure 4:**
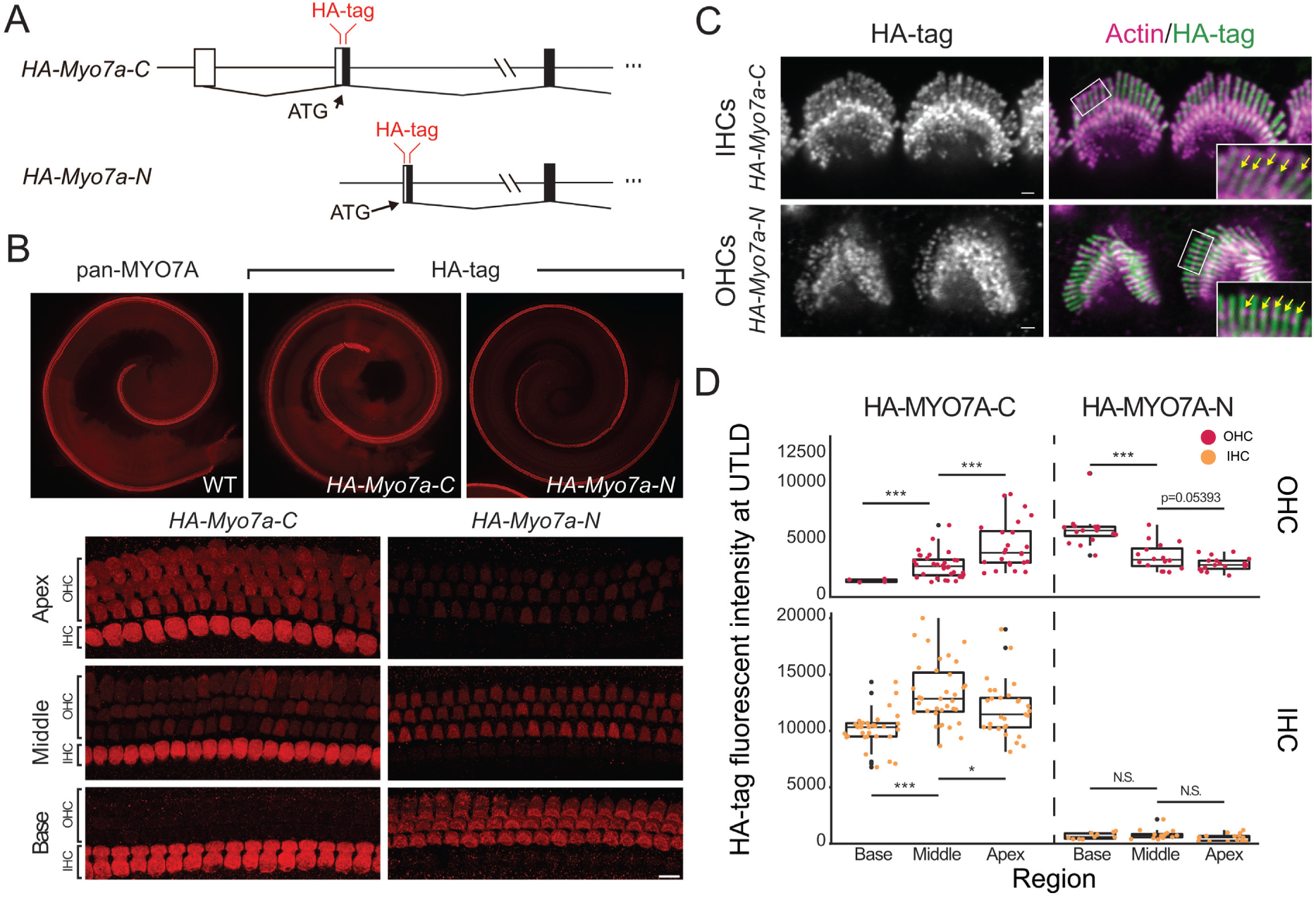
Both MYO7A-C and MYO7A-N localize to UTLDs of IHCs and OHCs. **A**. Diagram of the generation of *HA-Myo7a-C* and *HA-Myo7a-N* mice. HA-tag sequence is inserted after the translational start sites of *Myo7a-C* and *Myo7a-N*, respectively. **B**. Immunofluorescence imaging of whole mount WT type, *HA-Myo7a-C* and *HA-Myo7a-N* mice at P5. A pan-MYO7A antibody was used on WT cochlea to show the overall expression pattern of MYO7A. HA-tag specific antibody was used to visualize HA-MYO7A-C and HA-MYO7A-N in the KI mouse lines. (Scale bar=10μm) **C**. Immunofluorescence images showing that HA-signal is also detected in the UTLDs of IHCs and OHCs of both HA-MYO7A-C and HA-MYO7A-N mice. (Scale bar=1μm) **D**. Quantification of HA immunoreactivity at the UTLDs of *HA-Myo7a-C* and *HA-Myo7a-N* mice. Similar to their expression pattern in the cell body, the HA signal also showed tonotopic pattern in UTLDs of OHCs. (Statistics: Number of cells (cochlea): OHC: HA-MYO7A-C: base=4 (1) (4 of the cochlea have nearly non-detectable value, which is excluded from T-test), middle=32 (5), apex=24 (5); HA-MYO7A-N: base=16 (4), middle=32 (4), apex =24 (4); IHC: HA-MYO7A-C: base=32 (4), middle=35 (5), apex =29 (5); HA-MYO7A-N: base=8 (5) (1 of the cochlea have nearly non-detectable value, which is excluded from T-test), middle=16 (5), apex =15 (5). T-test p-value: OHC: HA-MYO7A-C: base vs middle= 4.06e-07, middle vs apex= 0.0007474; HA-MYO7A-N: base vs middle= 4.062e-05, middle vs apex= 0.05393; IHC: HA-MYO7A-C: base vs middle= 1.979e-07, middle vs apex= 0.03071; HA-MYO7A-N: base vs middle= 0.3507, middle vs apex= 0.05761). Box-plots show medians, 25th, and 75th percentiles as box limits and minima and maxima as whiskers.

We first investigated HA-MYO7A-C and HA-MYO7A-N levels in the P5 cochlea. As previously reported, HA-MYO7A-C was predominantly expressed in IHCs in all regions of the cochlea (**Fig. 4B**). HA-MYO7A-C is also relatively abundantly expressed in apical OHCs compared to the cochlear base. In contrast, HA-MYO7A-N was mainly expressed in OHCs, with higher levels in the basal region and decreasing toward the apex (**Fig. 4B**). This inversely correlated expression pattern of the HA-MYO7A-C and HA-MYO7A-N isoforms is consistent with the pattern we inferred from isoform-specific deletion mouse lines.

We next investigated the subcellular localization of MYO7A-N, especially at the functionally important site of the UTLDs. As previously shown for the HA-MYO7A-C mice^20^, we also detected HA-MYO7A-N immunoreactivity at the UTLD of OHCs (**Fig. 4C, Supplementary Fig. 2A**). Quantification analysis showed that HA-MYO7A-C is the predominant isoform in IHC UTLDs, consistent with its high expression level in the cell body (statistical analysis in **Fig. 4D** legend). We also observed a tonotopic gradient of MYO7A-C and MYO7A-N in UTLDs. OHCs have the highest HA-MYO7A-N/HA-MYO7A-C ratio in the basal region, decreasing toward the cochlear apex (**Fig. 4D**).

Given the proposed role of MYO7A in tip-link tensioning, we hypothesize that the tonotopically graded distribution of MYO7A-C and MYO7A-N within the UTLD of OHCs may contribute to the previously reported gradient of tip-link tension along the cochlear tonotopic axis^19^. This may occur through isoform-specific differences in motor structure and activity. We thus next investigated the relationship between the molecular structures and enzymatic functions of the motor domains of the two MYO7A isoforms.

### 3.6 Cryo-EM structure of the MYO7A motor domain and comparative analysis with other unconventional myosins

We performed high-resolution cryo-EM analysis on purified MYO7A-C-2IQ and MYO7A-N-2IQ isoforms, each containing the Myosin motor domain and two IQ domains, at the ADP-bound stage in complex with actin filaments. The resulting structures were resolved at 3.1 Å (MYO7A-C-2IQ) and 2.9 Å (MYO7A-N-2IQ) (**Fig. 5A**). To our knowledge, these are the first high-resolution structures of the mammalian MYO7A motor domain to be reported.

**Figure 5:**
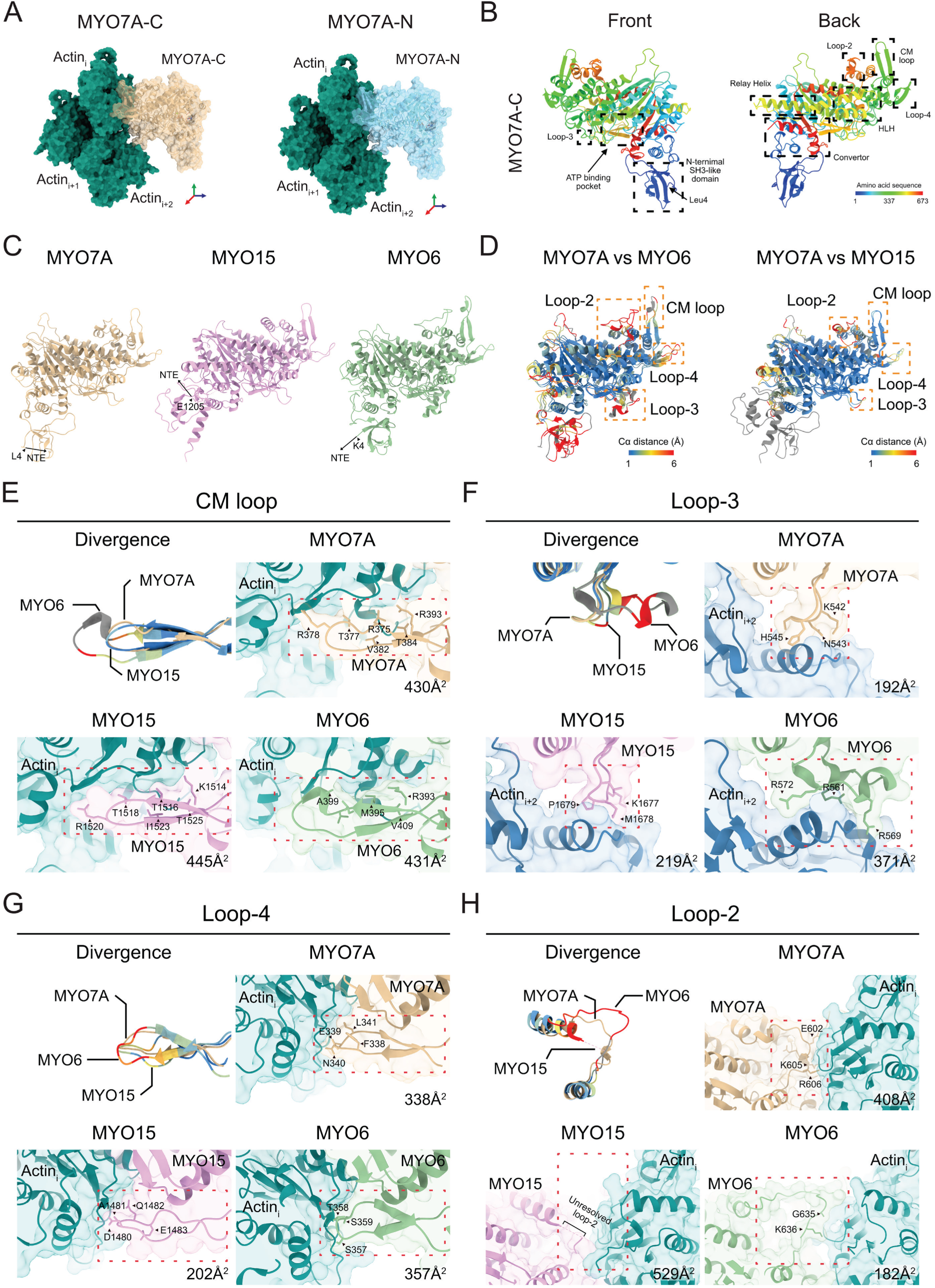
Structural comparison of MYO7A, MYO15 and MYO6. **A**. Cryo-EM structures of MYO7A-C (yellow) and MYO7A-N (blue) in complex with an actin filament bundle (green). For enhanced visualization, the structures are rotated 180° along the y-axis and 90° along the x-axis. **B**. Structural representation of MYO7A subdomains. The order of amino acid sequences are color-coded using a rainbow gradient. **C**. Overall structural models of MYO7A-C (yellow, UniProt Entry: P97479), MYO15 (pink, UniProt Entry: Q9QZZ4), and MYO6 (light green, UniProt Entry: Q64331). The first resolved residue for each model is labeled with arrowhead and the location of the unresolved NTE is labeled with arrow. (First resolved residues: MYO7A-C: Leu4; MYO15A: Glu1205; MYO6: Lys4). **D**. Divergence map highlighting differences between MYO7A-C and MYO15/MYO6. Actin-myosin interfaces where Cα distances exceed 6 Å are enclosed in yellow boxes. Residues are colored by blue-yellow-red gradient according to Cα distances. **E–H**. Enlarged views of key actin-myosin interfaces (CM loop, loop-3, loop-4, and loop-2). In the divergence panels, MYO15 and MYO6 are colored according to their Cα distance from MYO7A-C (yellow). Misaligned residues (due to additional or absent residues) compared with MYO7A-C are shown in gray. For each enlarged panel, corresponding actin-myosin interface is highlighted with red boxes. Key residues contributing to a high buried surface area are marked with arrowheads. Values of buried surface area of each actin-myosin interface are labeled in the bottom-right corner.

First, to investigate how the MYO7A ATPase differs from other members of the myosin superfamily, we performed a multi-sequence alignment and structural divergence analysis using previously reported cryo-EM structures of MYO15A^35^ and MYO6^36^ bound to ADP and actin. We found that the ATP-binding pockets, as measured by Cα–Cα distances and root mean square deviation of atomic positions (RMSD), remained highly conserved between these three myosins. A structural comparison of the ATP-binding pockets of MYO6, MYO15A and MYO7A, including the location of representative human mutations causing Usher syndrome 1B, is described in **Supplementary Fig. 3A, B**.

At the actomyosin interface, however, substantial structural divergence was observed between MYO6, MYO15A and MYO7A. To quantify the divergences in the actomyosin interface, we calculated the buried surface area, which reflects the extent of contact between proteins, and identified key residues mediating actin binding. Four structurally divergent loops, cardiomyopathy loop (CM loop), loop-2, loop-3, and loop-4 were analyzed for their contributions to actin binding^37^. The CM loop contributes to the strong binding state of myosin for actin^38–40^. Despite MYO6 having five additional residues, its buried surface area (431 Å²) is comparable to MYO7A (430 Å²) and MYO15A (445 Å²) (**Fig. 5E**). Loop-3 mediates interactions with actin_i+2_^41–43^. In MYO6, this loop contains nine additional amino acids, including three arginine residues, which significantly increase its buried surface area (317 Å²) compared to MYO7A (192 Å²) and MYO15A (219 Å²) (**Fig. 5F**). Loop-4 is known for its role in actin binding and tuning motor velocity^44–46^ and exhibited comparable buried surface areas for MYO6 (357 Å²) and MYO7A (338 Å²), while MYO15A displayed the smallest interface (202 Å²) (**Fig. 5G**). Loop-2 is suggested to modulate actin-activated ATPase activity^47–50^. Loop-2 of MYO15A contains ten additional residues. However, loop-2 of MYO15A was unresolved^35^, indicating it is dispensable for strong binding of MYO15A to F-actin (**Fig. 5H**). Despite MYO6 containing five additional residues compared to MYO7A, its buried surface area was significantly smaller (182 Å² vs. 408 Å²) (**Fig. 5H**). Overall, our structural analysis highlights sequence and structural divergence among unconventional myosins, particularly at the acto-myosin interface, which likely contribute to their kinetic functional specializations.

### 3.7 Allosteric changes in the MYO7A head domain induced by isoform-specific NTE domains

We next focused on potential differences in the molecular structure of the two MYO7A isoforms. Since MYO7A-C and MYO7A-N differ exclusively by their NTEs, we were particularly interested in the structure of these unique domains; however, both were unresolved in our cryo-EM derived models, indicating that the NTEs of both isoforms were likely unstructured in the ADP-bound, strong actin-binding state. We therefore explored the possibility that the distinct NTEs of the two isoforms might exert long-range allosteric effects on the structure of critical subdomains in the ATPase. While Root mean square deviation (RMSD) analysis revealed a high degree of overall structural similarity between the MYO7A-N and –C isoforms, with an RMSD of 1 Å (maximum Cα deviation of 7.3 Å) across 670 amino acids (**Fig. 6A, Supplementary Movie 1**), three regions exhibited significant local divergence, with deviations of Cα pairs exceeding 6 Å, including loop-2, the N-terminal SH3-like domain, and the relay domain. (**Fig. 6A, Supplementary Movie 1**)

**Figure 6:**
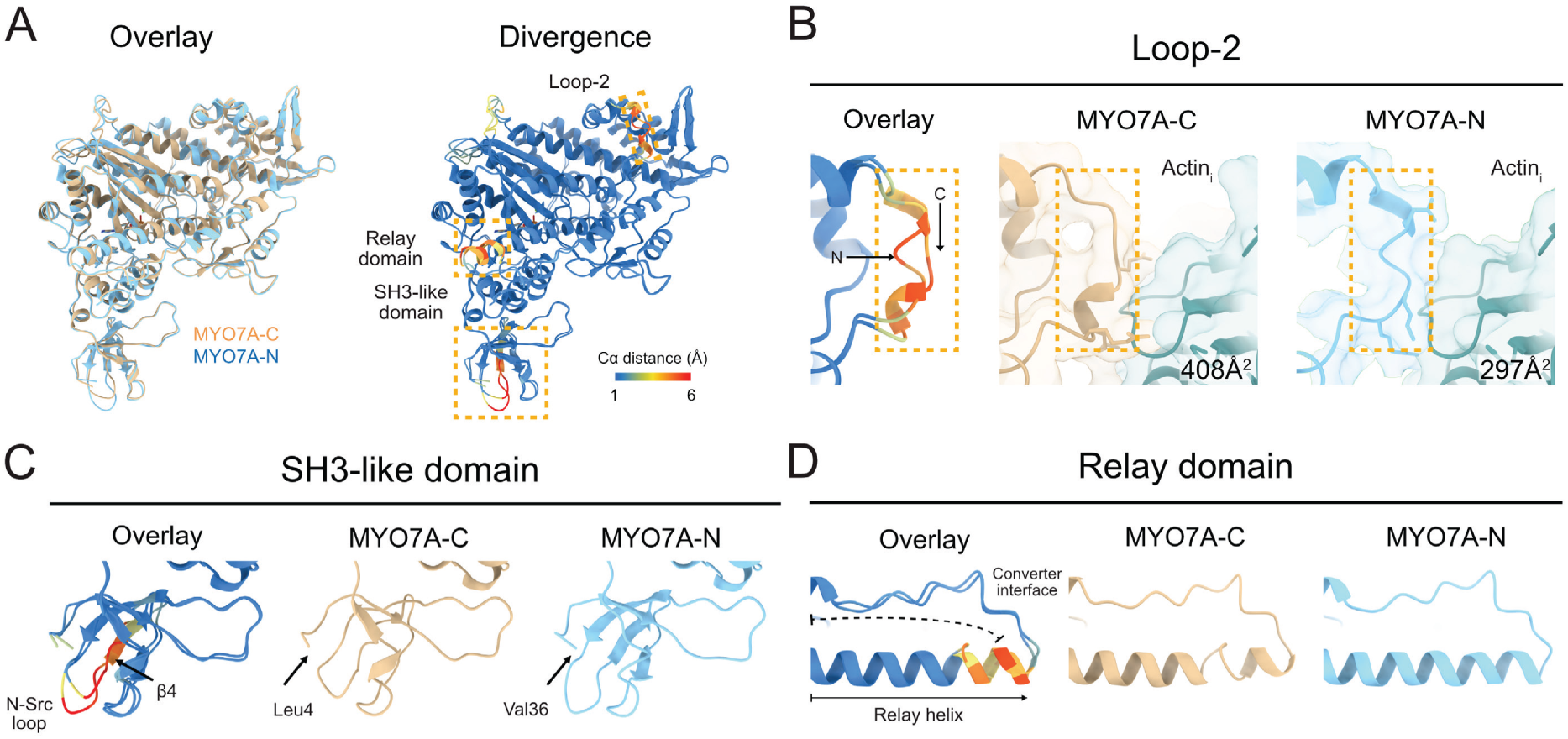
Structural differences between MYO7A-C and MYO7A-N. **A**. Divergence map of MYO7A-C and MYO7A-N subdomains with significant structural deviations (Cα distance > 6Å), including Loop-2, the SH3-like domain, and the relay domain, are highlighted within yellow boxes. **B**. Box-enlarged images of loop-2. MYO7A-C and MYO7A-N are indicated with arrows in the overlay panel. And the interactions between MYO7A-C/MYO7A-N with actin are shown in the right two panels, with MYO7A-C in yellow, MYO7A-N in blue and actin in green. **C**. Box-enlarged images of SH3-like domain. N-Src loop and β-sheet 4, which shows the highest Cα differences, are indicated in the overlay panel. The two arrows on the right two panels indicate the first resolved amino acid of MYO7A-C (Leu4) and MYO7A-N (Val36). **D**. Box-enlarged images of the relay domain. Arrow indicates the order of animo acid sequence, with N-terminus on the left and C-terminus of the relay domain on the right.

Loop-2, a critical subdomain involved in interactions with actin, is hypothesized to play a role in facilitating the initial weak stage myosin-actin binding and activating the ATPase^51–53^. Our cryo-EM analysis revealed notable structural divergence in loop-2 between MYO7A-C and MYO7A-N: 6 residues (MYO7A-C: G600-R606; MYO7A-N: G632-R638) showed RMSD of ∼6 Å, which is significantly higher than the overall RMSD (**Fig. 6B**). The deviation of these residues contributes to the differences of their buried surface areas with actin, with MYO7A-C showing a 408 Å² buried surface, compared to only 297 Å² for MYO7A-N (**Fig. 6B**).

The next notable structural divergence is located in the SH3-like domain, which is positioned between the NTE and the main motor domain (**Fig. 6C**). It has been implicated in modulating myosin activity through interactions with the myosin essential light chains (ELC)^54,55^ or autoinhibition^56^. We identified five residues of SH3-like domain (MYO7A-C: S33-V38; MYO7A-N: S65-V70) with significant RMSD ∼6 Å (**Fig. 6C**). These five residues form part of the N-src loop and the third β-sheet of the SH3-link domain. Additionally, MYO7A-C exhibited a likely disrupted of β2 and β4-sheet of SH3-like domain in the model (**Fig. 6C**). Considering the high vicinity of the SH3-like domain to the NTE, these structural alterations may result from the confirmation influence of NTEs. Potentially, this divergence could impact interactions with the ELC or regulatory light chain (RLC) of MYO7A, thereby potentially influencing motor functionality.

We also identified potential structural variances in five residues on the C-terminus of the relay helix (MYO7A-C: E475-E480; MYO7A-N: E507-E512), with RMSD ∼6 Å (**Fig. 6D**). The relay helix links confirmational changes from motor domain to the converter and lever arm^57^. Our structure analysis suggests that the α-helix structure at the C-terminus of the MYO7A-C relay helix is likely to be disrupted in our model (**Fig. 6D**). Although the converter domain is not resolved in our cryo-EM data, it is shown to interact with the C-terminus of the relay helix in other species of myosins. This divergence of the C-terminus of the relay helix between MYO7A-C and MYO7A-N could potentially affect the confirmational linkage between the motor domain and converter.

### 3.8 MYO7A-C and MYO7A-N have distinct ATPase enzymatic properties

We next investigated whether the distinct NTEs, and the subtle but significant structural differences in loop-2, SH3-like and relay domains could affect the enzymatic and mechanic activities of MYO7A-C and MYO7A-N at the UTLD. To test this hypothesis, we measured the steady-state, actin-activated ATPase activity of purified MYO7A-C-2IQ (C-2IQ), and separately MYO7A-N-2IQ (N-2IQ) (**Fig. 7A**) using the NADH-coupled assay at 25°C under saturating ATP conditions. By titrating increasing amounts of F-actin into the reaction, the catalytic rate (*k_cat_*) of N-2IQ (*k_cat_* = 0.05 ± 0.003 s^-1^) was significantly reduced compared with C-2IQ (*k_cat_* = 0.38 ± 0.02 s^-1^) (**Fig. 7D, E**). The apparent affinity for F-actin (*K_ATPase_*), representing the concentration of F-actin required to reach half-maximal ATPase activation, was not significantly different between C-2IQ (*K_ATPase_* = 2.7 ± 0.5 μM) and N-2IQ (1.4 ± 0.6 μM) (**Fig. 7D, F**). Basal ATPase rates (without F-actin) were unchanged between N-2IQ (*k_basal_* = 0.013 ± 0.003 s^-1^) and C-2IQ (*k_basal_* = 0.028 ± 0.002 s^-1^). We conclude that in comparison to MYO7A-C, the altered NTE of MYO7A-N reduced the maximal catalytic activity of the ATPase without altering its apparent affinity for F-actin (Table 1).

**Figure 7:**
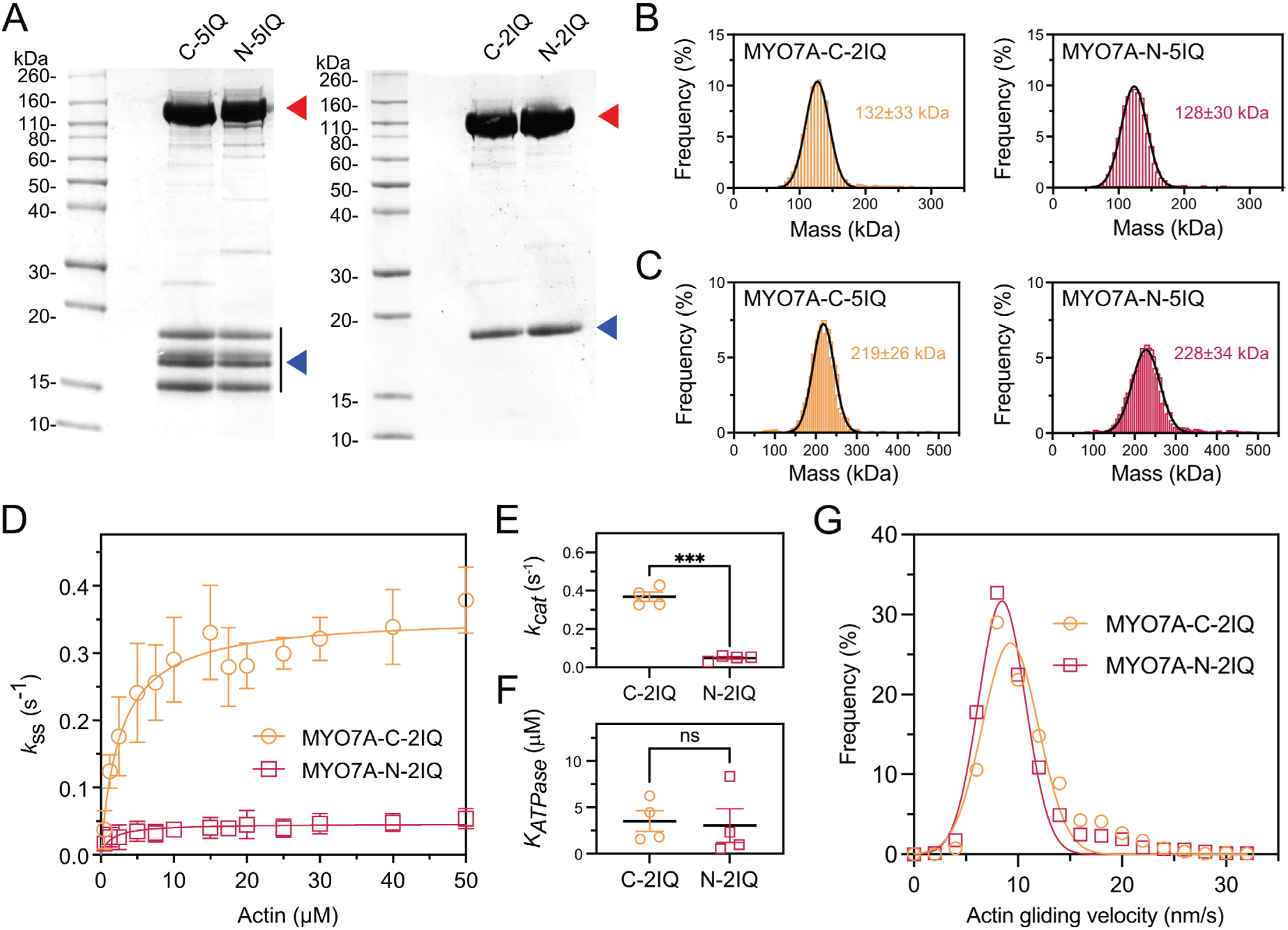
The enzymatic activity of MYO7A isoforms is differentially tuned. **A.** SDS-PAGE analyses of purified MYO7A-C and MYO7A-N motor domain truncations used in this study. The ATPase from mouse MYO7A-C, or –N, was truncated after the 2^nd^ or 5^th^ IQ light-chain binding site and expressed in Sf9 cells. The myosin heavy chain (∼110 kDa) (red arrow) co-purified with light chains (LCs) ranging from 15 - 20 kDa (blue arrow). C-2IQ / N-2IQ co-purifies with MYL12B (RLC), and C-5IQ / N-5IQ co-purifies with MYL12B, MYL6, CALM and CALML4. **B**. Single molecule measurements of purified MYO7A 2IQ truncations using mass photometry (MP). Average weights of 132±33 kDa (C-2IQ) and 128±30 kDa (N-2IQ) were estimated by Gaussian fitting. **C.** Single molecule MP measurement of MYO7A 5IQ truncations. Average weights of 219±26 kDa (C-2IQ) and 228±34 kDa (N-2IQ) were estimated by Gaussian fitting. N=7333-8389 particles from 2 myosin prep. **D.** Measurement of actin-activated ATPase activity by the NADH steady-state assay at 25°C. Hyperbolic fits to averaged data estimates the maximum catalytic rate (*k*_cat_) and [actin] to reach half maximal activation (*K*_ATPase_) for C-2IQ (0.38±0.02 s^-1^, 2.7±0.5 μM, blue line), and N-2IQ (0.05±0.003 s^-1^, 1.4±0.6 μM, red line). Data points are mean ± SD from 4 independent determinations and 2 myosin preps. **E+F.** Statistical analysis of *k*_cat_ and *K*_ATPase_ from hyperbolic fits to individual titrations. Identical data reanalyzed from (D). ***, P< 0.001, n.s., non-significance, unpaired t-test with Welch’s correction. **G.** Measurement of myosin mechanical activity using the *in vitro* gliding filament assay at 30°C. Gaussian fits are shown for C-2IQ (blue, 9.3 ± 2.7 nm/s) and N-2IQ (red, 8.5 ± 2.3 nm/s). N = 1173-2020 filaments analyzed from 2 myosin prep.

**Table 1.** Buried surface area of MYO7A-actin interfaces. Buried surface area between actin binding subdomains of MYO7A isoforms and actins (Å²).

|  |  | Overall | CM<br>loop | Loop 4 | Loop-2 | HLH | Loop 3 |
| --- | --- | --- | --- | --- | --- | --- | --- |
| MYO7A-C | Actin i | 1199 | 376 | 126 | 408 | 306 | 0 |
|  | Actin i+2 | 311 | 0 | 0 | 0 | 153 | 164 |
| MYO7A-N | Actin i | 1311 | 444 | 175 | 297 | 363 | 0 |
|  | Actin i+2 | 365 | 0 | 0 | 0 | 201 | 158 |

**Table 2.** Parameters for the steady-state ATPase assay.

|  | **MYO7A-C-2IQ** | **MYO7A-N-2IQ** |
| --- | --- | --- |
| $k_{\text{cat}} (\text{s}^{-1})$ | $0.36 \pm 0.01$ | $0.05 \pm 0.003$ |
| $K_{\text{ATPase}} (\mu\text{M})$ | $2.7 \pm 0.5$ | $1.4 \pm 0.6$ |

Light chains binding to MYO7A motor domain can regulate its ATPase activity^58,59^. We therefore investigated whether this phenomenon might be responsible for the measured differences in ATPase activity between isoforms. MYO7A has five IQ domains that are reported to be a binding site for calmodulin (CALM), regulatory light chain (RLC; MYL12B) and calmodulin-like protein 4 (CALML4)^59–63^. We co-expressed N-2IQ and C-2IQ with CALM, MYL12B, and CALML4, in addition to essential light chain (ELC, MYL6) that has been reported as a light chain for other non-muscle myosins^64^. Both N-2IQ and C-2IQ co-purified with a single light chain analyzed by SDS-PAGE (Fig. 7A), and for both isoforms this was unambiguously identified as MYL12B using LC-MS/MS. Purified N-2IQ and C-2IQ holoenzymes were further analyzed by mass photometry, a label-free optical technique to weigh single molecules^65^. Mass photometry yielded an average molecular weight of 132 ± 33 kDa for C-2IQ, consistent with the ATPase (119 kDa) plus a single RLC (20 kDa), and 126 ± 30 kDa for MYO7A-N-2IQ, also consistent with the ATPase (122 kDa) plus a single MYL12B (20 kDa) (**Fig. 7B**). We extended this light chain analysis to MYO7A-N and MYO7A-C proteins truncated to retain all 5IQs (N-5IQ and C-5IQ). Both proteins co-purified with 4 distinct low molecular weight light chains, and these were identified as CALM, MYL12B, MYL6 and CALML4 using LC-MS/MS (**Fig. 7A**). Mass photometry measured average molecular weights of 228 ± 34 kDa (N-5IQ) and 219 ± 26 kDa (C-5IQ) (**Fig. 7C**). These results were in close agreement with predicted molecular weights of 216 kDa (N-5IQ) and 213 kDa (C-5IQ), calculated assuming an equal stoichiometry of 1:1:1:1 of MYL6, MYL12B, CALM, CALML4 binding to each ATPase. We conclude there was no intrinsic difference in light chain binding between truncated MYO7A-C and MYO7A-N molecules, and that this was unlikely the cause of altered ATPase activity.

To estimate the relative mechanical output of MYO7A-C vs MYO7A-N isoforms, we used an *in vitro* gliding filament assay^66^. Imaging chambers were functionalized with either purified C-2IQ or N-2IQ protein, and rhodamine-labelled actin filaments were bound to the myosin-functionalized surface. The velocity of actin filaments, propelled across the surface by myosin activity, was recorded using TIRF microscopy in the presence of saturating ATP at 30°C (**Supplementary Movie 2, 3**). The average gliding velocities were 9.3 ± 2.7 nm•s-1 (C-2IQ) and 8.5 ± 2.3•s-1 (N-2IQ), indicating a modest reduction in the mechanical output of N-2IQ compared with C-2IQ (**Fig. 7G**). In conclusion, our results show differing ATPase activities of MYO7A isoforms, with MYO7A-N having a significantly reduced catalytic rate compared to MYO7A-C.

## 4. Discussion

The molecular properties of MYO7A have been well characterized, particularly in the context of Usher syndrome type 1B^22,23,67^. Its robust and widespread expression in hair cells has made it a standard marker, yet its primary functional role has remained somewhat elusive. At the stereocilia base, MYO7A forms a multiprotein complex with PCDH15, and in the context of mechanotransduction, contributes, alongside SANS and Harmonin, to the upper tip-link density^18,20^.

We previously demonstrated that auditory hair cells express multiple MYO7A isoforms. The canonical isoform, MYO7A-C, is enriched in IHCs and expressed in OHCs along a tonotopic gradient that decreases toward the cochlear base. Loss of MYO7A-C delays MET current onset and reduces resting open probability in IHCs, supporting a role in tip-link tensioning^20^.

In the present study, we identify MYO7A-N, a complementary isoform predominantly expressed in OHCs, with a reciprocal tonotopic pattern—rising from apex to base. We also examined MYO7A-S, a variant lacking the N-terminal extension (NTE), and found it to be expressed at very low levels, and to have minimal functional relevance in mature hair cells. Collectively, these findings define MYO7A-C and MYO7A-N as the principal isoforms in auditory hair cells, each with distinct spatial expression and likely specialized functions.

### Functional consequences of MYO7A isoform loss

Our data show that loss of MYO7A isoforms leads to distinct hearing phenotypes. *Myo7a-ΔC* mice show severe ABR deficits with preserved DPOAEs, consistent with MYO7A-C’s role in IHCs. In contrast, *Myo7a-ΔN* mice exhibit progressive, milder hearing loss and reduced DPOAEs, aligning with MYO7A-N’s role in OHCs and cochlear amplification. MYO7A-N expression is highest in basal cochlear regions; thus, more severe high-frequency hearing loss is expected beyond the 32 kHz limit of our current ABR assays.

### Structural and enzymatic differences between MYO7A isoforms

Previous structural insights into MYO7A have primarily relied on molecular dynamics simulations^58^. In this study, we present high-resolution cryo-EM models of the two mouse MYO7A isoforms. Comparative analysis of MYO7A with MYO6 and MYO15 revealed notable differences at the actin-myosin interface, which likely underlies their distinct biochemical properties. The ATP-binding pockets of these three myosins on the other hand exhibit relatively minor divergence. This is consistent with previous findings that these myosins all are considered high duty-ratio myosins. Overall, our structural models provide opportunities for further detailed comparisons of unconventional myosins important for hair cell function.

Although the NTEs of both MYO7A isoforms were unresolved in cryo-EM, likely due to intrinsic disorder, their presence appears to modulate distal structural features and motor activity. Between MYO7A-C and MYO7A-N, we identified three subdomains that exhibit significant structural divergences. Among them, loop-2 deserves special attention due to its well-established role in actin-activated ATP hydrolysis activity^47–50^. Loop-2 in MYO7A-C shows greater buried surface area with actin correlating with higher ATPase activity in the enzymatic assays. The ATPase activity results suggest MYO7A-C undergoes more rapid actin detachment-reattachment cycles, whereas MYO7A-N may spend more time in a strong actin-bound state^68^. We thus propose that MYO7A-N is equipped with a superior ability to generate and maintain mechanical tension compared to MYO7A-C. While we have not performed enzymatic studies with the short isoform MYO7A-S, a recent study compared enzymatic activities of MYO7A-C and MYO7A-S and found that the ATPase activity of MYO7A-S was lower than MYO7A-C^59^, comparable to the value we determined for MYO7A-N in the present study. Although the detailed biochemical properties of all MYO7A isoforms remain to be more fully explored, our findings support the idea that the NTEs of MYO7A isoforms influence their motor activity, potentially giving rise to functional differences in hair cells. This is not unique for MYO7A: There is strong precedent for NTEs modulating myosin activities and cellular functions. For instance, a 35-aa NTE of MYO1C significantly reduces its velocity and force output in actin gliding assays^69,70^. Similarly, different MYO15A isoforms are localized to different rows of hair bundles, regulating stereocilia length^71^.

### A potential role of NTEs in phase separation?

Disordered regions like the NTE of MYO7A-N are commonly implicated in liquid–liquid phase separation (LLPS), forming dynamic, membraneless condensates that concentrate specific protein complexes^72^. LLPS is emerging as a mechanism for organizing the MET machinery in stereocilia. Key proteins, including Whirlin, Harmonin, SANS, PCDH15, and CDH23, have been hypothesized to form condensates critical for UTLD and ankle-link complex assembly^73–76^. Given the disordered nature of MYO7A-N’s NTE and its known binding partners, we hypothesize that MYO7A-N may participate in stereocilia condensates, potentially influencing spatial organization or scaffolding of MET complexes. Thus, in addition to enzymatic differences, MYO7A isoforms may diverge in their capacity to engage in LLPS, providing a new layer of regulation.

### Potential roles of MYO7A isoforms in the auditory system

A central question raised by our findings is why distinct MYO7A isoforms, each with different motor kinetics, are differentially expressed across hair cell types and along the tonotopic axis. Resting open probability, considered a proxy for tip-link tension, differs significantly between hair cell populations, with IHCs exhibiting ∼10–20% Po and OHCs ∼50%^19^. Additionally, tip-link tension increases from apex to base in OHCs^19^, paralleling the gradient of MYO7A-N expression. These correlations suggest that MYO7A isoforms might contribute to spatial tuning of MET tension, potentially optimizing hair cell function across frequencies.

Two models may explain the mechanistic role of MYO7A isoforms in this context: a “dynamic motor” model and a “static anchor” model. The dynamic motor model, historically proposed to explain tip-link tension generation^3,33,77,78^, directly aligns with our current and previous findings^20^. It posits that myosin molecules actively contribute to tip-link tension. In this framework, multiple MYO7A molecules interact with the tip link or scaffold proteins at the UTLD, and their dynamic upward movement generates tip-link tension. This model is supported by evidence that MYO7A functions as a processive motor in various organisms^59,61,79,80^. Our structural and biochemical data suggest MYO7A-C hydrolyzes ATP more rapidly, enabling quicker detachment-reattachment cycling, while MYO7A-N binds actin more tightly, potentially generating sustained force. The ratio of MYO7A-C to MYO7A-N at the UTLD could therefore modulate tension. This could contribute to frequency tuning in OHCs, serving as a molecular frequency filter in addition to the tuning provided by basilar membrane mechanics and other mechanisms.

Alternatively, the static anchor model proposes that MYO7A serves primarily as a structural anchor rather than a force generator. In this view, MYO7A isoforms do not actively pull on tip links but instead just stabilize their connections to actin filaments. The MYO7A-C to MYO7A-N ratio could modulate actin-binding affinity, allowing the UTLD to accommodate varying resting tensions set by other mechanisms, such as MET-dependent cytoskeletal remodeling or changes in membrane tension. This model aligns with recent findings showing that MYO7A is essential for tip-link stability^24^.

These models are not mutually exclusive. MYO7A may function as a motor during developmental stages to establish tension, then transition into an anchor role to maintain it—similar to MYO6’s behavior under load^81^. While the motor and anchor models provide plausible mechanisms for how MYO7A isoforms regulate tip-link tension, these hypotheses remain speculative and require further experimental validation.

### Implications for Gene Therapy

Our findings also have clear implications for inner ear gene therapy. Current approaches typically target the canonical MYO7A isoform. However, the differential expression and distinct functional roles of MYO7A-C and MYO7A-N suggest that isoform-specific replacement may be necessary for optimal therapeutic outcomes. A nuanced understanding of isoform biology is essential for developing effective treatments for MYO7A-related hearing loss.

## Figures

**Supplementary figure 1:**
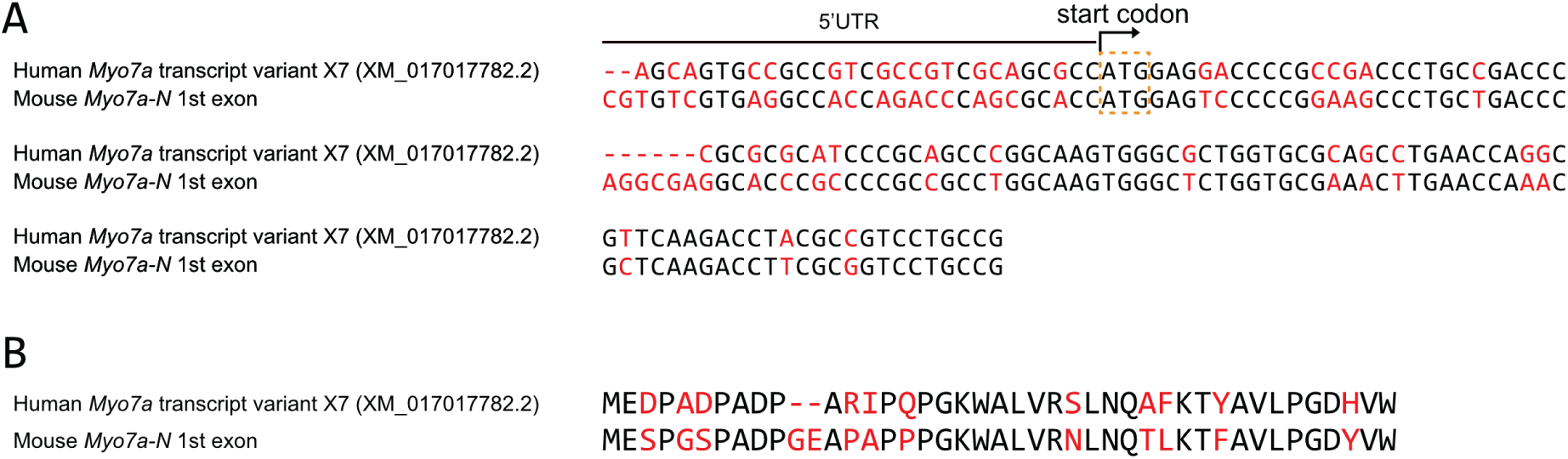
Conservation between human and mouse MYO7A-N. **A**. Alignment of human MYO7A-N (PREDICTED: Homo sapiens myosin VIIA (MYO7A), transcript variant X7, mRNA, XM_017017782.2) and mouse *Myo7a-N* first exon nucleotide sequence. The mismatched bases are highlighted with red. **B**. Alignment of human *MYO7A-N* (translation of XM_017017782.2) and mouse MYO7A-N NTEs amino-acid sequence. The mismatched bases are highlighted with red.

**Supplementary figure 2:**
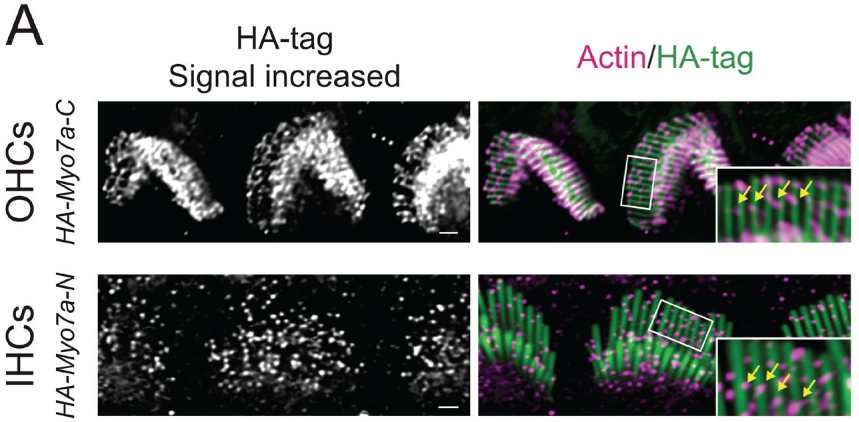
visualization of UTLDs in MYO7A-C OHCs and MYO7A-N IHCs. **A**. Immunofluorescence images showing that HA-signal is also detected in the UTLDs of HA-MYO7A-C OHCs and HA-MYO7A-N IHCs. (Scale bar=1μm)

**Supplementary figure 3:**
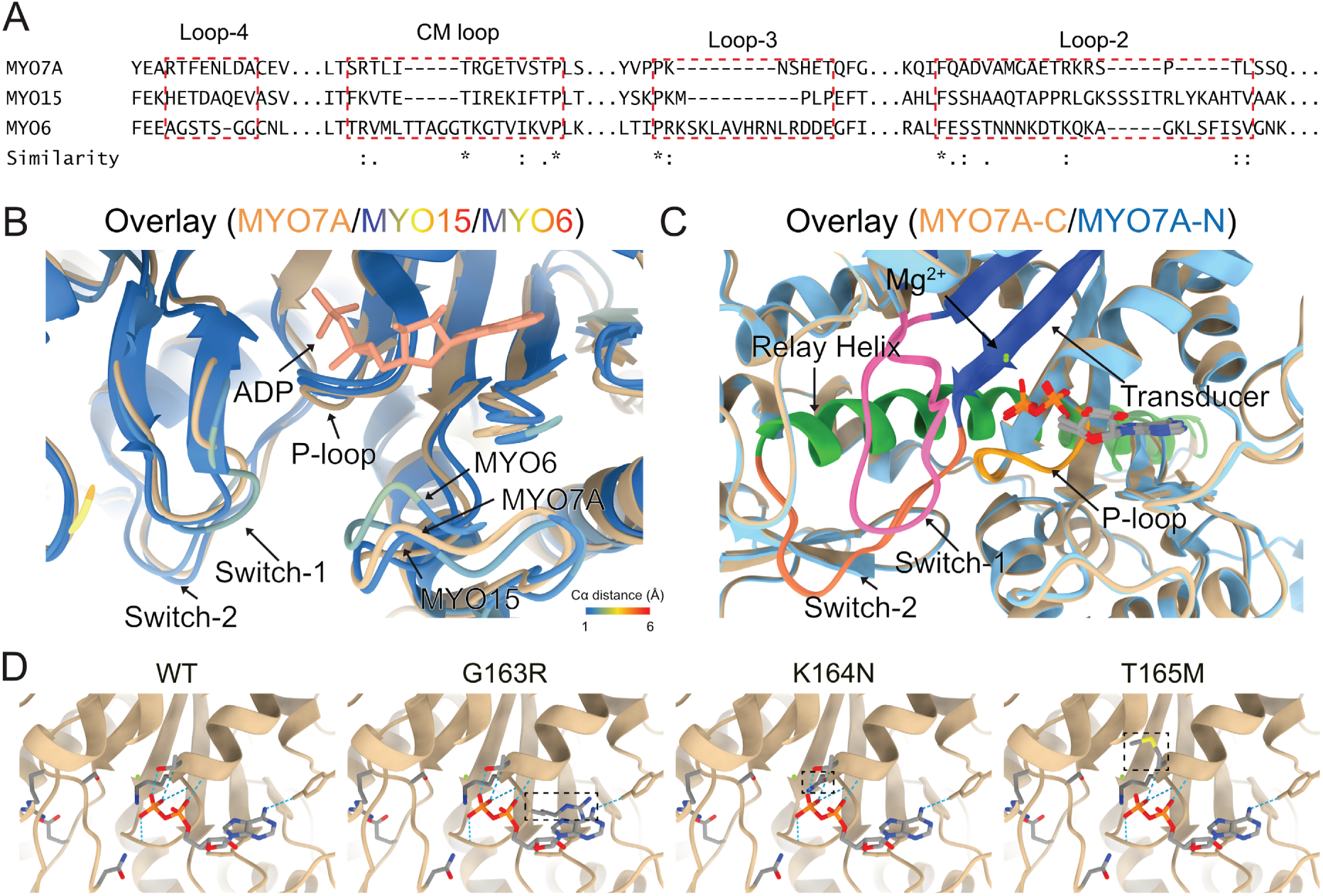
Structural comparison of MYO7A, MYO15A, and MYO6 ATP-binding pocket. Structure of ATP-binding pockets among MYO7A, MYO15A, and MYO6 are relatively similar compared to their divergence in the actomyosin interfaces. We next characterized critical subdomains within the MYO7A ATP-binding pocket, including Switch-1, Switch-2, and the P-loop. We also examined how deafness-associated mutations impact the structure and function of MYO7A’s ATP-binding pocket. Specifically, we analyzed three Usher syndrome-related mutations — G163R^82–84^, K164N^85,86^, and T165M^82,85,87^ — to assess their structural effects. Compared to previous molecular dynamics models, our cryo-EM structure provides a more precise and reliable prediction of these mutations’ impact. G163R substitutes a neutral glycine with a bulky arginine, potentially sterically hindering ATP binding and impairing hydrolysis efficiency. K164N replaces lysine with asparagine, with the potential to disrupt H-bonds within the P-loop and reduce its stability during the powerstroke. T165M substitutes threonine with methionine, disrupting H-bonds with Mg²⁺. It also disrupts the coordination and hydrolysis of ATP, likely to impair MYO7A’s motor function and contribute to Usher syndrome pathology. **A**. Multiple sequence alignment between MYO7A, MYO15 and MYO6. Actin-myosin interfaces (loop-4, CM loop, loop-3 and loop-2) are highlighted with red boxes. **B**. Divergence map of ATP-binding pocket of MYO7A, MYO15 and MYO6. MYO7A is shown in yellow and MYO15/MYO6 is colored by their Cα distance to MYO7A. Color scale is shown in the bottom-right corner. **C**. Detailed structure of the ATP-binding pockets in MYO7A. The transducer, P-loop, relay helix, switch 1, and switch 2 regions are labeled in dark blue, yellow, green, pink, and orange, respectively. **D**. ATP-binding pocket showing the mapped human deafness-associated mutations (T163R, K164N, T165M). Models generated based on the cryo-EM structure of MYO7A-C.

## 5. Method

### 5.1 Animal care and handling

The care and use of animals for all experiments described conformed to NIH guidelines. Experimental mice were housed in a 12:12 hour light:dark cycle with free access to chow and water in standard laboratory cages in a temperature and humidity-controlled vivarium. The protocols for the care and use of animals were approved by the Institutional Animal Care and Use Committees at the University of Virginia, the University of Colorado Denver, and the National Institute on Deafness and Other Communication Disorders. All the institutions mentioned above are accredited by the American Association for the Accreditation of Laboratory Animal Care. C57BL/6J (Bl6, from Jackson Laboratory, ME, USA) mice and sibling mice served as control mice for this study. Neonatal mouse pups (P0–P5) were sacrificed by rapid decapitation. Mature mice were euthanized by CO_2_ asphyxiation followed by cervical dislocation.

### 5.2 Cochlea single-cell long-read RNA Sequencing

The sample preparation protocol was adapted from a previous publication by the Kelley lab. The organs of Corti were dissected from approximately ten postnatal day 7 (P7) mice of both sexes in cold Dulbecco’s Modified Eagle Medium (DMEM, Thermo Fisher Scientific, MA, USA). The collected samples were incubated with 0.25% trypsin-EDTA (Thermo Fisher Scientific, MA, USA) at 37°C for 10 minutes. To quench trypsin activity, an equal volume of fetal bovine serum (FBS) was added. The samples were then gently triturated 40 times on ice, followed by filtration through a 40 µm cell strainer. Single cells were collected by centrifugation at 300 × g and resuspended in phosphate-buffered saline (PBS, Thermo Fisher Scientific, MA, USA) containing 0.04% FBS. Cell concentration and viability were assessed using an automated cell counter (RWD-C100, RWD Life Science, TX, USA). Samples with a cell concentration of 800–1000 cells/µL and a viability of >95% were used for sequencing. Single-cell cDNA library preparation was performed at the Genome Analysis and Technology Core, University of Virginia. The EpiMultiome ATAC + Gene Expression kit (10x Genomics, CA, USA) was used to generate single-cell cDNA libraries, targeting a recovery of 10,000 cells. For long-read sequencing, 75 ng of the single-cell cDNA library was used as the recommended input for MAS-ISO-seq library preparation. The MAS-ISO-seq kit (102-407-900, Pacific Biosciences, CA, USA) was used to generate concatemers for long-read sequencing. Library quality was assessed using a Qubit fluorometer (Q33238, Thermo Fisher Scientific, MA, USA) and an Agilent 2100 Bioanalyzer (G2939BA, Agilent Technologies, CA, USA). The final MAS-ISO-seq library was submitted to Maryland Genomics (MD, USA) for sequencing on the PacBio Revio system (102-090-600, Pacific Biosciences, CA, USA). Raw sequencing files were processed using the PacBio Iso-Seq package (version 4.2.0, https://isoseq.how/). Processed reads were mapped to the mouse genome (GRCm39, M36). Transcript classification was performed using Pigeon (version 1.3.0, https://isoseq.how/classification/pigeon.html). Single-cell analysis was conducted using the Seurat package (version 5.2.0, https://satijalab.org/seurat/). All bioinformatics analyses and computations were performed on the University of Virginia High-Performance Computing systems.

### 5.3 5’ Rapid Amplification of cDNA Ends (RACE)

5’RACE is conducted to validate the transcriptional start sites of *Myo7a* isoforms. Total RNA was extracted using TRIzol reagent (15596026, ThermoFisher Scientific) from ∼10 P7 animals. Then the RACE is performed using 5’ RACE System for Rapid Amplification of cDNA Ends, version 2.0 (18374058, ThermoFisher Scientific) according to the protocol. *E. coli* cDNA library containing *Myo7a* transcripts are cloned by TA Cloning™ Kit (K202020, ThermoFisher Scientific). *E. coli* colonies are selected by *Myo7a* isoforms specific primers. Positive colonies are then sent for whole plasmid Sanger sequencing (Eurofins genomics DNA Sequencing Lab, KY, USA) to identify the sequence of *Myo7a* transcripts.

### 5.4 Generation of Myo7a-ΔC, Myo7a-ΔN, Myo7a-ΔS, Myo7a-ΔCN, Myo7a full KO, HA-Myo7a-N, and HA-Myo7a-C mice

For CRISPR/Cas mediated generation of the mouse models, we used the online tool CRISPR (http://crispor.tefor.net/crispor.py) to select suitable target sequences to ablate Myo7a isoforms. To generate the corresponding single-guide (sg)RNAs, a PCR product from overlapping oligonucleotides (as described in the CRISPR online tool) was generated by T7 in vitro transcription. In vitro transcription was performed using the MAXIscript T7 kit (Life Technologies), and RNA was purified using the MEGAclear kit (Thermo Fisher Scientific, Waltham, MA). To produce genetically engineered mice, fertilized eggs were coinjected with Cas9 protein (PNA Bio, 50 ng/μl) and the sgRNA (30 ng/μl). Two-cell stage embryos were implanted on the following day into the oviducts of pseudopregnant ICR female mice (Envigo). Genotyping was performed by PCR amplification of the region of interest. Target sequences, sgRNAs, repair templates, and genotyping primers are listed below:

*Myo7a-ΔC* and *HA-Myo7a-C* mice:

sgRNA: AAGCATGGTTATTCTGCAGA**AGG**, targeting the second exon of *Myo7a-C* (1^st^ coding exon).

Repair template for *HA-Myo7a-C* mice:

GCCTGGGCTCAGGGCGTGCCATGGTCTCTTCCCACAGAGCTGTGTCTGGTCACTCCGGCAGGTGTGCTGACGTAGAAGCATGTACCCATACGATGTTCCAGATTACGCTGTTATTCTGCAAAAGGTGAGTGCGTCTCCTCTCTCTCAGAGCTGCAGAGGGCCATGCTGGGTACCTCACATCCCACCCT GCA.

Genotyping primers: GCTCTGGTCACATAGACTGAGCT (forward), and GGGATGAATCATCACTCCTGC (reverse).

*Myo7a-ΔN* and *HA-Myo7a-N* mice:

sgRNA: AGGGCTTCCGGGGGACTCCA**TGG**. This sgRNA targets the first coding exon of *Myo7a-N* (exon1).

*HA-Myo7a-N* Repair template:

GTTTAGGGTTCCCAAGCGGGTGGCGTGCGGGATTAGGTGAATTAGGAGCCCGCTTCGTGTCGTGAGGCCACCAGACCCAGCGCACCATGTACCCATACGATGTTCCAGATTACGCTGAGTCCCCCGGAAGCCCTGCTGACCCAGGCGAGGCACCCGCCCCGCCGCCTGGCAAGTGGGCTCTGGTGCGAAACTTGAACCAA

Genotyping primers: GTGCTCCCTCCCCTACAGTA (forward), CTTTGGTGTGAACTTGGCCG (reverse).

*Myo7a-ΔCN* mice:

sgRNA: CAAACTCCTGGCCTGACTTC**AGG**, targeting the common exon of *Myo7a-C* (exon 3), *Myo7a-N* (exon 2), and *Myo7a-S* (exon 2).

Repair template for *Myo7a-ΔCN* mice: we deleted one thymine in the reading frame of *Myo7a-C* and *Myo7a-N*. This T deletion is in the untranslated region of *Myo7a-S*. GGTAGTGGGAGAGAGGGGCTCAACTTCAGGATCTCCTGCCTACAGGGGGACTATGATGGATGGACCTGAAGTCAGGCCAGGAGTTTGATGTGCCCATCGGGGCCGTGGTGAAGCTCTGCGAC.

Genotyping primers: TTGGTGTGCCTGCCTTAGT (forward), and ACCTGCCATTGTAGCCTTCC (reverse).

*Myo7a full KO*:

sgRNA: GCCTGATGAAGACGAGGAGGACC. This sgRNA created a 26bp deletion in exon 24

Genotyping primer: CCAGCCTAACGGTTAAGACA (forward), AGCTGGTCACCCTCATCGT (reverse).

*Myo7a-ΔS* mice:

We designed two sgRNAs that flank the entire first exon of *Myo7a-S*. This method created two DNA double-strand breaks and deleted 275bp, which covers the first exon of *Myo7a-S* (this exon is 107bp).

sgRNAs: CAGCCCTGCCTAAGACAGTG**TGG** (beginning of the exon), GGGGTGTAGTTTGTGGGGTC**AGG** (end of the exon).

Genotyping primers: CAGAGCCTATCTATAGCCTTGTGC (forward), AAGGATGACCTCCCTTCCAT (reverse).

*Myo7a-ΔEnhancerA* mice:

sgRNAs: ACAACAGCGACTTCCCTGCA**TGG** and GCGAGACTGGCTAGGAACGT**AGG**. Similar strategy with *Myo7a-ΔS* mice, we created an approximately 400bp deletion that covers the entire *EnhancerA*.

Genotyping primers: ACCATGACCCCATAAGGTTC (forward), AGTGTTCTCCCGAGAGCCTTTC (Reverse).

### 5.5 Quantitative polymerase chain reaction (qPCR)

qPCR primers were designed to detect either *pan-Myo7a* transcripts or *Myo7a* isoform-specific transcripts to evaluate their expression levels. Total RNA was extracted using TRIzol reagent (15596026, ThermoFisher Scientific). Reverse transcription was performed using the SuperScript™ IV Reverse Transcription system (18090010, ThermoFisher Scientific). qPCR reactions were conducted using iTaq Universal SYBR Green Supermix (1725121, Bio-Rad, CA, USA) on a CFX Opus 384 Real-Time PCR System (12011319, Bio-Rad) for fluorescence measurement. Following analysis and plotting is done by R (version 4.4.1). Primers:

### 5.6 Immunofluorescence

Inner ear organs were fixed in 3% paraformaldehyde (PFA, Electron Microscopy Sciences, PA) immediately after dissection for 20 min. Samples were washed three times with phosphate-buffered saline (GIBCO^®^ PBS, Thermo Fisher Scientific, Waltham, MA) for 5 min each. After blocking for 2 h with blocking buffer (1% bovine serum albumin, 3% normal donkey serum, and 0.2% saponin in PBS), tissues were incubated in blocking buffer containing primary antibody at 4 °C overnight. The following antibodies were used in this study: rabbit polyclonal Myosin-VIIa antibody (catalog#: 25-6790, Proteus Biosciences Inc, Ramona, CA. 1:100), mouse monoclonal Myosin-VIIa antibody (Developmental Studies Hybridoma Bank, MYO7A 138-1, concentrate, 1:100), mouse monoclonal Myosin-VI antibody (A-9, Santa Cruz, 1:100), rabbit anti-Harmonin antibody (H3, obtained from Ulrich Mueller’s lab), rabbit anti-ADGRV1 antibody (obtained from Dominic Cosgrove’s lab), rabbit polyclonal Myosin-VIIa antibody used in Fig. 2 (PB206) was custom-generated and affinity purified against the immunizing MYO7A peptide LPGQEGQAPSGFEDLERGR, and rabbit anti-HA antibody (C29F4, Cell Signaling Technologies, Danvers, MA). Fluorescence imaging was performed using a Leica Stellaris 5 Confocal Microscope Platform or Zeiss LSM880 with AiryScan.

### 5.7 Hearing tests in mice

ABRs of WT and *Myo7a-ΔC* mice at P17, 4-week, 6-week, and 9-week were determined. Mice were anesthetized with a single intraperitoneal injection of 100 mg/kg ketamine hydrochloride (Fort Dodge Animal Health) and 10 mg/kg xylazine hydrochloride (Lloyd Laboratories). ABR and DPOAE were performed in a sound-attenuating cubicle (Med-Associates, product number: ENV-022MD-WF), and mice were kept on a Deltaphase isothermal heating pad (Braintree Scientific) to maintain body temperature. ABR recording equipment was purchased from Intelligent Hearing Systems (Miami, Fl). Recordings were captured by subdermal needle electrodes (FE-7; Grass Technologies). The non-inverting electrode was placed at the vertex of the midline, the inverting electrode over the mastoid of the right ear, and the ground electrode on the upper thigh. Stimulus tones (pure tones) were presented at a rate of 21.1/s through a high-frequency transducer (Intelligent Hearing Systems). Responses were filtered at 300–3000 Hz, and threshold levels were determined from 1024 stimulus presentations at 8, 11.3, 16, 22.4, and 32 kHz. Stimulus intensity was decreased in 5–10 dB steps until a response waveform could no longer be identified. Stimulus intensity was then increased in 5 dB steps until a waveform could again be identified. If a waveform could not be identified at the maximum output of the transducer, a value of 5 dB was added to the maximum output as the threshold.

DPOAEs of the same group of WT and *Myo7a-ΔC* mice were recorded. While under anesthesia for ABR testing, DPOAE was recorded using SmartOAE ver. 5.20 (Intelligent Hearing Systems). A range of pure tones from 8 to 32 kHz (16 sweeps) was used to obtain the DPOAE for the right ear. DPOAE recordings were made for *f*2 frequencies from 8.8 to 35.3 kHz using a paradigm set as follows: *L*1 = 65 dB, *L*2 = 55 dB SPL, and f_2_/f_1_ = 1.22.

### 5.8 Scanning electron microscopy

Adult mice were euthanized by CO2 asphyxiation before intracardiac perfusion with 2.5% glutaraldehyde (Electron Microscopy Sciences, Hatfield, PA) and 2% PFA. The otic capsule was dissected and incubated in postfixation buffer at 4 °C overnight (2.5% glutaraldehyde, in 0.1 M cacodylate buffer, with 3 mM CaCl2). For neonatal mouse pups, the samples were dissected and treated with postfixation buffer immediately. The otic capsules from adult mice were incubated for two weeks in 4.13% EDTA for decalcification and then further dissected to expose the organ of Corti. Samples underwent the OTOTO procedure and were dehydrated using gradient ethanol and critical point drying. After sputter coating with platinum, the samples were imaged on Zeiss Sigma VP HD field emission SEM using the in-lens secondary electron detector.

### 5.9 Actin purification and labeling

Actin was extracted from rabbit skeletal acetone powder (Pel-Freeze) in G-buffer (2 mM Tris-HCl pH 8, 0.2 mM NaATP, 0.1 mM CaCl2, 1 mM NaN3, and 0.5 mM DTT) and purified by cycles of polymerization and depolymerization, as described^88^. F-actin was dialyzed extensively against 4mM MOPS pH 7.0, 2 mM MgCl2 0.1 mM EGTA, 3 mM NaN3, 1 mM DTT prior to use. The concentration of F-actin was determined by measuring the absorbance at 290 nm (ε = 26,600 M−1 cm−1). For preparation of fluorescently labeled actin filaments, F-actin was incubated with equimolar rhodamine-phalloidin (Invitrogen) for 30 mins before use, in a buffer of 20 mM MOPS pH7.5, 5 mM MgCl2, 0.1 mM EGTA, 100 mM KCl, and 1 mM DTT.

### 5.10 Cryo-EM Sample preparation and data collection

Lacey grids were glow-discharged for 20s using a GloQube. 3µL of F-actin at 0.5µM concentration was applied to a lacey grid, incubated for 5s, then aspirated. Next, 3µL of the F-actin was applied to the grid, then 3µL of myosin at 3µM was applied to the grid and incubated for 5s. 3µL was aspirated and the rest of sample was back blotted for 3s at 99% humidity using a Leica EM GP. Grids were flash-frozen in liquid ethane.

Data collection was done at 300keV on a Titan Krios (Thermo Fisher Scientific) equipped with a K3 direct electron detector (Gatan). 40-frame movies were collected at 1.08 Å/px with total fluence of 50e^-^/A^2^. On-the-fly data preprocessing and quality control were performed using cryoSPARC Live. Motion correction and CTF estimation were done in cryoSPARC (ver. 4.3.1).

Regions of F-actin with myosin bound were manually selected from several micrographs to generate 2D class averages used for automatic template-based particle picking by Template Picker in cryoSPARC. Particles were extracted with box size 512x512px and subjected to several rounds of 2D classification. Homogenous refinement job was done using particles containing F-actin with bound myosin and a volume generated during cryoSPARC Live session. Per-particle defocus refinement was done using Local CTF Refinement job, and then homogenous refinement was run again. Discrete heterogenous states of F-actin:MYO7A-C-2IQ were analyzed by 3D Classification in cryoSPARC using particles with orientations and shifts from homogenous refinement job, a mask focused on the myosin motor domain, and four classes as input. The resulting volumes were visually inspected and particles from a class with well-defined continuous density for the myosin motor domain were taken for another round of homogenous refinement. To further increase signal from the myosin motor domain the Local Refinement job was performed using particles and volume from homogenous refinement and a mask focused on the N-terminal portion of the myosin motor domain. The output volume was post-processed by EM Ready.

An initial MYO7A-C-2IQ model was generated by AlphaFold, and rebuilt to fit the density obtained from the 3D reconstruction. The model was refined in Coot and Phenix using the density obtained from homogenous refinement before post-processing by EM Ready.

Cryo-EM analysis, including calculation of RMSD, buried surface area, mutation modeling and visual representation was performed using UCSF ChimeraX. Subdomain identifications were based on previous publications and conservation analysis by using the BLAT tool from Uniport Database (https://www.uniprot.org/).

### 5.11 Mass photometry

The mass of purified MYO7A-C/N holoenzymes was measured using label-free mass photometry (OneMP, Refeyn). High tolerance #1.5 coverslips (Thorlabs, 24 x 50 mm) were cleaned by isopropanol and ddH2O with sonication for 5 min each, then dried under a stream of filtered nitrogen. A silicon gasket was placed on the cleaned coverslips to create sample chambers of ∼2 mm diameter. MYO7A holoenzymes were diluted to a final concentration of 50 nM in 10 mM MOPS pH 7.0, 250 mM KCl, 0.15 mM EGTA, and 1 mM NaN3, and added to the sample chamber. Single molecule landing events were collected for 90 seconds using AcquireMP (Refeyn), and data were filtered and processed using DiscoverMP (Refeyn). The mass distribution of single MYO7A molecules was fitted with a Gaussian curve using Prism (GraphPad v10).

### 5.12 Steady-state ATPase assay

Steady-state ATPase activity of the MYO7A holoenzyme (200 nM) was measured using a NADH enzyme-linked assay in a final buffer of 10 mM MOPS pH 7.0, 25 mM KCl, 2 mM MgCl2, 0.15 mM EGTA, 2 mM MgATP, 200 uM NADH, 40 units/mL lactate dehydrogenase, 200 units/mL pyruvate kinase and 1 mM phosphoenolpyruvate. The timecourse of NADH absorbance was measured at 340 nm using a dual-beam spectrophotometer (UV-1800, Shimadzu) with a multicell cuvette thermostatically held at 25°C. The rate of ATP consumption was calculated from gradient of 340 nm absorbance. Each measured rate was corrected for basal ATPase activity in the absence of F-actin, and for the intrinsic ATPase activity of F-actin. ATP hydrolysis rates were fit to the Michaelis-Menten equation to estimate kcat and KATPase using Prism (GraphPad v10).

### 5.13 In vitro actin gliding assay

Clean cover glass (#1.5, 24 x 50 mm) was coated with 0.1% nitrocellulose in amyl acetate (Ladd Industries) and attached to a glass slide using two strips of double-sided adhesive tape spaced 3 mm apart to form an imaging chamber. The chamber was incubated for 5 mins with a GFP antibody (GFP-20, G6539, Sigma-Aldrich), blocked with 1 mg/mL BSA (A0281, Sigma-Aldrich), and washed with 20 mM MOPS pH 7.5, 5 mM MgCl2, 0.1 mM EGTA, 100 mM KCl, 1 mM DTT. Chambers were incubated with purified MYO7A holoenzyme (0.3 mg/mL) for 1 minute and washed. To block any non-functional myosin heads, chambers were then incubated with 2 uM phalloidin stabilized F-actin that was shredded through a fine-gauge needle to generate small filaments. After a 1 minute incubation, chambers were washed with wash buffer supplemented with 2 mM ATP. Chambers were next incubated with rhodamine-labeled F-actin (100 nM) for 1 minute, and then washes into motility buffer (20 mM MOPS pH 7.5, 5 mM MgCl2, 0.1 mM EGTA, 150 mM KCl, 2 mM ATP, 50 mM DTT, 3 mM glucose, 100 ug/mL glucose oxidase, and 20 ug/mL catalase) at 30°C. Filaments were visualized using total internal reflectance fluorescence (TIRF) microscopy (Nikon H-TIRF) using a 100x oil objective (N.A 1.49, Nikon Apo TIRF) and an inverted microscope stand (Ti2-E). Time-lapse images were captured on an EM-CCD camera (Andor iXon DU-897) controlled by NIS-Elements (AR version 5.42.06, Nikon). Images were processed in FiJi^89^ and stage drift removed using the Stabilizer plugin. Filament velocities were calculated using TrackMate^90^.

## Acknowledgments

We thank Katia Sol-Church from Genome Analysis and Technology Core (University of Virginia, RRID:SCR_018883) for generating single-cell RNA sequencing library. We thank Gloria Sheynkman and Natchanon Sittipongpittaya at University of Virginia for generating long-read sequencing library and support of data analysis. We thank Luke Tallon from Maryland Genomics (University of Maryland) for PACBIO long-read sequencing. We thank Wenhao Xu and Daniel Grigsby at Genetically Engineered Murine Model Core (University of Virginia, RRID:SCR_025473) for generating mouse models used in this study. We thank Sijie Hao from Advanced Microscopy Facility (University of Virginia, RRID: SCR_018736) for the support of SEM imaging. We thank Ewa Niedzialkowska from Egelman lab for performing cryo-EM imaging. We thank Molecular Electron Microscopy Core (University of Virginia, RRID:SCR_019031) for providing services of cryo-EM imaging.

## Author contributions

Conceptualization, S.L., J.P., J.E.B., J.B.Shin; methodology, S.L., J.P., J.E.B., E.H.E., J.B. Shin; experiments, S.L., J.P., T.M.P., J.E.B., E.H.E., J.B. Shin; writing—review & editing, S.L., J.P., J.E.B., E.H.E., J.B. Shin; funding acquisition, J.E.B., E.H.E, J.B. Shin.; supervision, J.E.B., E.H.E., J.B. Shin.

